# Impacts of global warming on fertility of male fishes: evidence from meta-analysis of temperature effects on spermatozoa motility

**DOI:** 10.1101/2024.04.24.590878

**Authors:** Yu Cheng, Jinhai Wang, Sayyed Mohammad Hadi Alavi, Songpei Zhang, Zuzana Linhartová, Deepali Rahi Roy, Nururshopa Eskander Shazada, Borys Dzyuba, Otomar Linhart

**Affiliations:** University of South Bohemia in České Budějovice, Faculty of Fisheries and Protection of Waters, South Bohemian Research Center of Aquaculture and Biodiversity of Hydrocenoses, Research Institute of Fish Culture and Hydrobiology, Zátiší 728/II, Vodňany 389 25, Czech Republic; School of Fisheries, Aquaculture and Aquatic Sciences, Auburn University, 203 Swingle Hall, Auburn, AL 36849, USA; School of Biology, College of Science, University of Tehran, Enqhelab Avenue, Tehran 14176-14411, Iran

**Keywords:** Fish reproduction, Global warming, Spermatozoa energy, Spermatozoa motility traits, Male fishes, Meta-analysis

## Abstract

Studies have demonstrated adverse effects of global warming on aquatic ecosystems from the abiotic to the biotic level. In the present work, a meta-analysis study was conducted to elucidate the effects of global warming on spermatozoa functions, which are key determinants of male fertility. We recruited 245 data records from a pool of empirical studies, which includes 20 studies spanning 20 cold- and warm-water fish species, to identify the effects of increased water temperature (IWT) on determinants of sperm fertility in fishes. Data were systematically re-processed and re-analyzed to determine the overall effects of IWT on sperm kinetics such as motility (MOT), duration of motility (DSM), curvilinear velocity (VCL), rectilinear velocity (VSL) and average path velocity (VAP), as well as on enzymatic activities for energy supply (EAES) and antioxidant enzyme activity (ANEA). The standardized mean difference was calculated for each study, with positive values indicating higher performance under IWT. Results showed that (a) the overall effect size for MOT was more negative in cold-water fishes (-1.22) than in warm-water fishes (-0.98). (b) Each 1 °C increase in the activation medium reduced MOT by 1.30% (cold-water fishes) and 3.47% (warm-water fishes). (c) The IWT negatively affected DSM, decreasing it by 10 s (cold-water species) and 5.64 s (warm-water species) per degree of IWT. (d) Spermatozoa velocity (VCL and VSL) was increased by IWT in warm-water species. In conclusion, this study shows that IWT negatively affects sperm motility kinetics, suggesting an impact of global warming on fish reproduction.

## 1. INTRODUCTION

Global changes in surface temperatures related to climate change are occurring at an unprecedented speed leading to multifarious significant impacts on both freshwater and marine ecosystems.^1^ It has been estimated that the oceans absorb over 93% of extra heat from the greenhouse effect.^2^ In this regard, Maberly et al.^3^ predicted a temperature rise by as much as 4°C for freshwater lakes by 2100. There are >30,000 fish species (www.fishbase.org) that are mostly thermotropic and possess an external fertilization strategy. Therefore, regulation and integration of various systems in their body including the reproductive system are directly affected by environmental temperature. In fishes, gamete maturation and spawning occur in a relatively narrow temperature range at which spermatozoa motility is activated to fertilize the oocytes.^4^ Current knowledge demonstrates the negative impacts of global temperature rise on fish biodiversity,^5,6^ on the sex ratio within wild populations,^7^ on reproduction, development and growth.^8,9^ These may affect sustainable fisheries and aquaculture.^10^ It has been hypothesized that thermally driven changes to physiological regulation of reproduction results in poor quality and quantity of gametes.^11–13^ Also, changes in environmental temperature are usually associated with advancement, delay or inhibition of spawning (gamete release) that subsequently affect reproductive success and survival of larvae in their natural environment. In fish reproduction, motility of spermatozoa is essential for fertilization and is triggered after discharge from their body into the aquatic environment where spermatozoa meet oocytes released from females.^14^ Identification and characterization of factors that affect spermatozoa motility provide significant basic and applied information to manage artificial reproduction in a hatchery practice or to predict reproductive success in wild populations. So far, several physiological and environmental factors have been identified that affect spermatozoa motility kinetics including temperature, pH, ions and osmolality (see reviews).^15–19^ Among them, the effects of temperature and its detrimental effects on spermatozoa motility kinetics remain largely unknown.

Spermatozoa motility kinetics are determinants of fertility in male fishes (see reviews)^20–22^ that include percentage of motile spermatozoa (MOT), duration of spermatozoa motility (DSM) and spermatozoa velocity such as curvilinear velocity (VCL; the actual velocity between its first and last detected positions), straight line velocity (VSL; the time-average velocity of a sperm head along the straight line between its first and last detected positions), and average path velocity (VAP: velocity along a smoothed path). Spermatozoa motility kinetics can be assessed using computer-assisted sperm analysis (CASA) from a video of spermatozoa motility recorded from an optical microscope.

It has been shown that a changing environmental temperature affects spermatozoa motility kinetics associated with the alternation of intracellular ionic concentrations or with the induction of oxidative stress.^23–29^ The impacts of environmental temperature on spermatozoa motility and velocity are largely ambiguous in fishes, displaying high variations between cold-water and warm-water fishes that naturally spawn in cold and warm water aquatic environments. Moreover, temperature effects on spermatozoa motility kinetics may vary within cold-water or warm-water fish species exhibiting species specificity. For example, Vladiĉ and Jätrvi^30^ found no differences in DSM when brown trout (*Salmo trutta*) spermatozoa were activated at 2–28 °C, while increasing temperature decreased DSM in Atlantic salmon (*Salmo salar*). Lahnsteiner and Mansour^31^ reported the highest spermatozoa motility and velocities for brown trout and burbot (*Lota lota*) at 4–6 °C and for grayling (*Thymallus thymallus*) at 8–16 °C. More recently, Merino et al.^19^ performed a systematic review on the effects of elevated temperature of the activation medium on spermatozoa motility kinetics in fishes. The inconsistency of findings across studies that might be due to experimental designs or protocols, methods for analysis of spermatozoa motility or species specificity, make it still difficult to interpret whether global warming affects spermatozoa motility functions with subsequent impacts on fish reproduction. Also, the estimated effects of increased temperature on spermatozoa motility kinetics are very heterogeneous probably due to the species investigated. Some studies have examined temperature ranges that are not ecologically or physiologically relevant or similar spermatozoa motility kinetics have not been assessed. Thus, a meta-analysis and synthesis of existing evidence based on optimal spawning temperatures are essential to draw reliable conclusions.

In the present study, we explored how elevated temperatures influence sperm quality through multiple moderator analyses such as on fish species based on our initial metadata in cold- and warm-water fishes. Meta-analysis is a technique used to collect and statistically evaluate results from numerous individual studies that explore similar research questions, leading to comprehensive and broader interpretations.^32^ Our main objective was to quantitatively synthesize empirical data, which examined the effects of increased water temperature (IWT) on spermatozoa motility kinetics including MOT, DSM, VCL, VSL and VAP in cold-water and warm-water fishes. To better understand the temperature effects, enzymatic activities for energy supply (EAES) and antioxidant enzyme activity (ANEA) were characterized. Our results provide novel information to elucidate the effects of increased temperature on spermatozoa motility kinetics with consideration of species specificity. It was also to determine the environmental temperature at which a particular species spawns, and the key molecular mechanisms involved in spermatozoa response to a temperature above their normal range.

## 2. MATERIALS AND METHODS

### 2.1. Literature search

The literature search and data collection were performed according to the Preferred Reporting Items for Systematic Reviews and Meta-Analyses (PRISMA) guidelines.^33^

The PRISMA flowchart for the search strategy and exclusion procedure is shown in Figure 1. Literature searches were performed in the databases for science including Web of Science, PubMed, Scopus and other sources such as Google Scholar and Citations. Articles in the English language published in peer-reviewed journals by 20 January 2024 were searched. The first search was adjusted to the following terms: (“temperature”) AND (“sperm” OR “spermatozoa” OR “spermatozoon” OR “semen” OR “male gametes”) AND (“motility rate” OR “motility duration” OR “velocity” OR “DNA fragment” OR “flagellum” OR “ATP consumption” OR “metabolism”) AND (“aquaculture” OR “aquatic animals” OR “fish” OR “marine”) AND (“fertilization”). Results of our search resulted in finding 166 papers that have examined the effects of temperature on spermatozoa motility kinetics in 24 fish species. Fourteen species were cold-water fishes with a preferred reproductive temperature at 4-15 °C and 10 species were warm-water fishes with a preferred reproductive temperature >16 °C (Figure 2). The classification of warm-water and cold-water fishes was according to FishBase (www.fishbase.se). When studies were screened for eligibility, 20 articles covering 20 fish species with a total of 245 data lines were used in the present study (Supplementary S1). The eligibility criterion examined was that temperatures had to be in the range of the naturnal spawning temperature (Figure 1).

**Figure 1.**
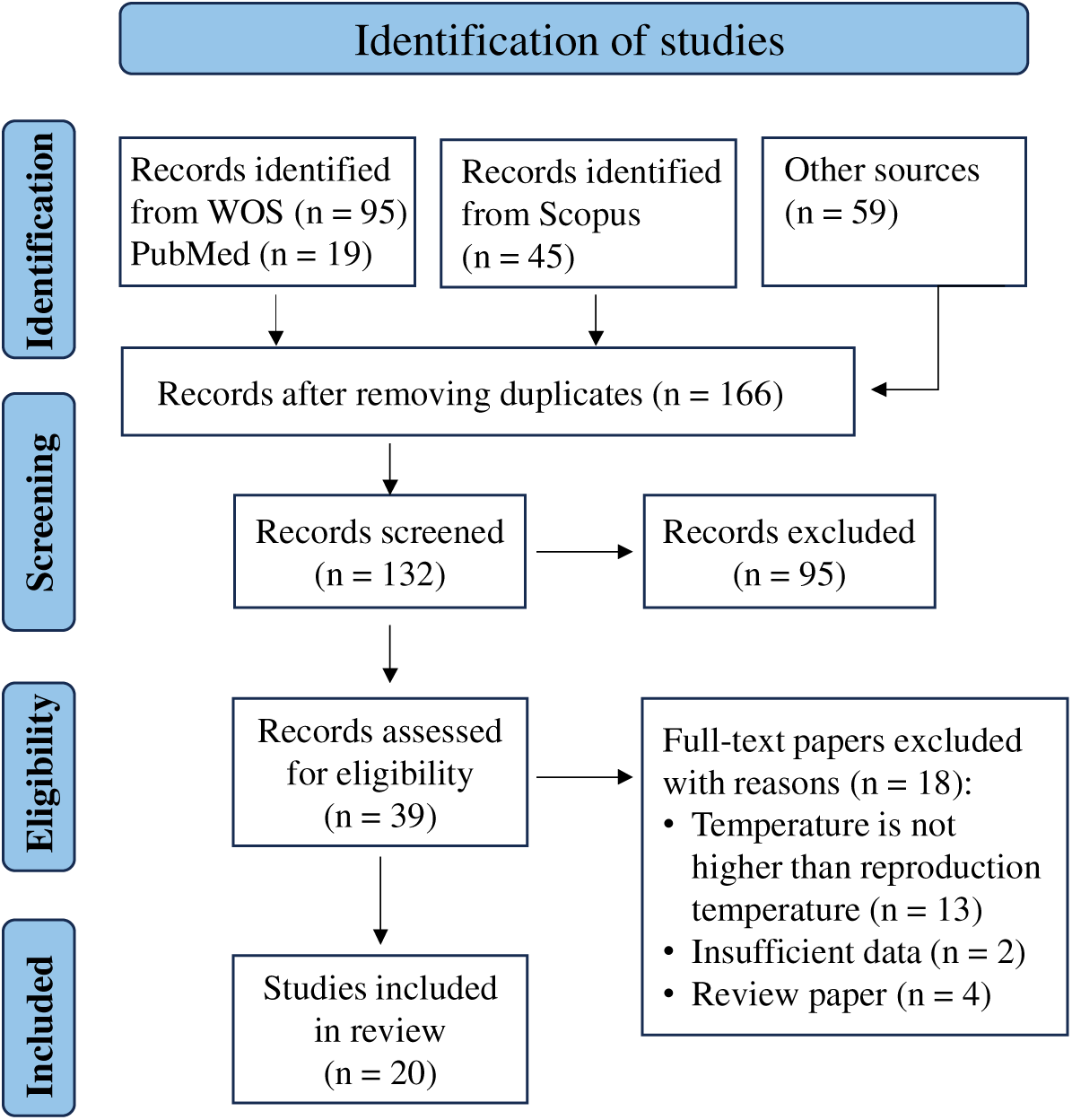
Preferred reporting items for systematic reviews and meta-analyses (PRISMA) flowchart used to search literature and to collect the data for the meta-analysis of temperature effects on spermatozoa motility kinetics in fishes. Numbers in parentheses show the number of studies.

**Figure 2.**
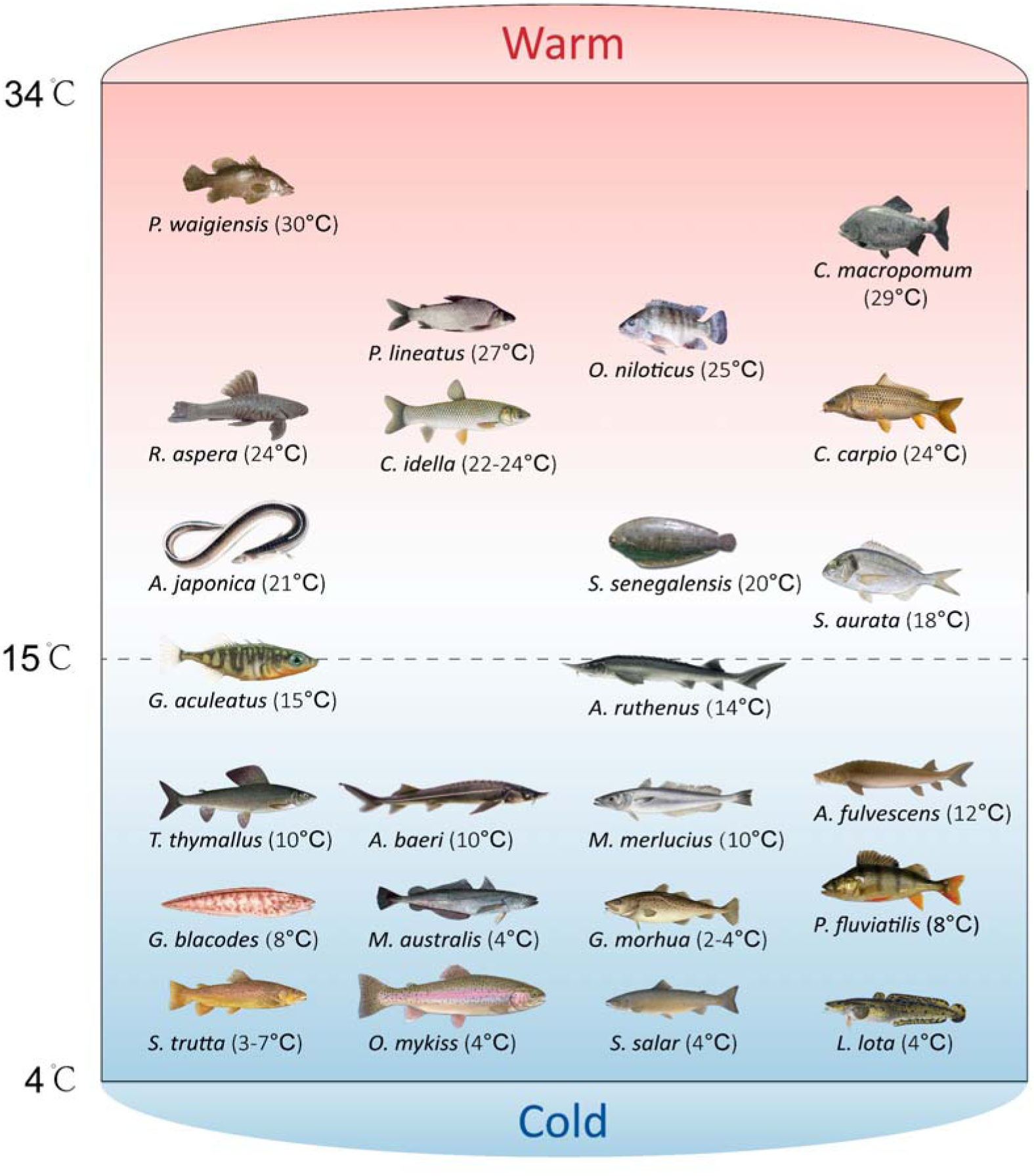
A diagram showing fish species that the influence of temperature on spermatozoa motility kinetics, enzyme activity and antioxidant response has been investigated. Temperatures in parentheses represent spawning temperature under natural conditions.

### 2.2. Criteria for study selection and data collection

The criteria considered to collect data for meta-analysis were: moderators of meta-analysis including (1) fish species, (2) time post-activation at which spermatozoa motility kinetics were assessed, (3) types of enzymes for energy supply and antioxidants, and (4) specific value and application scope for temperature. Only articles that examined different temperatures of the activation medium for spermatozoa activation and provided the specific value of temperature were selected. The temperature at which the fish species spawn naturally was considered as a control. All these parameters had to be reported in both control and elevated temperature groups. For a study that did not report standard deviations (S.D.) or standard errors of mean (S.E.), although these values could be inferred from existing data, we automatically filled them in using the imputation method.^34^

The following standards were considered to exclude data in meta-data analysis: (1) temperature of the activation medium was not increased. Of note, studies that exposed males to different temperatures were also excluded. (2) Temperature of the activation medium was increased for motility activation of short-term stored sperm. (3) Temperature of the activation medium was not specified. For instance, Lahnsteiner^35^ reported a range of temperature for MOT and spermatozoa, while actual (or exact) value of temperature was not specified. These kinds of data were not within our scope of investigation in the context of the temperature moderator. (4) Temperatures examined were lower than the species natural spawning temperature. For instance, Dadras et al.^26^ investigated the sensitivity of carp (*Cyprinus carpio*) spermatozoa to temperatures, however, the temperatures examined (4 and 14 °C) were lower than 24 °C that males release the sperm naturally. (5) Extraction of valid data from figures of the articles was not possible. For instance, Castro et al.^27^ reported the effects of temperature on enzyme and oxygen consumption rates using boxplots without data points causing difficulty in extracting mean values.

### 2.3. Extraction of data

From each study selected, the following information was extracted: author information, year of publication, title of article, fish species, time post sperm activation, enzyme type and sample size. Means and S.D. of MOT, DSM, VCL, VSL, VAP, EAES and ANEA for spermatozoa in the control and elevated temperature groups were extracted. When S.E. were reported, S.D. were calculated using the formula S.D. = S.E. × (*n*)^1/2^, where *n* is sample size. The open ImageJ software (https://imagej.nih.gov/ij/) was also used to extract data (mean, S.D. or S.E.) when they were shown in the figures of the articles.

### 2.4. Analyzing data

Statistical analyses were performed using the “metafor”^36^ and “orchaRd” packages^37^ in RStudio (version 4.2.2, 11 January 2022). The significance level for all statistical tests was set at *P* < 0.05.

#### Effect size

In a meta-analysis, the results of different trials must be combined using effect sizes that summarize the sign and magnitude of the effect reported in each trial.^38^ The spermatozoa motility kinetics were considered as continuous variables, and the Hedges’ g was used to calculate effect sizes.^38^

To compare the effects of the control and IWT on each variable, random-effects models were used to estimate overall mean effect sizes with 95% confidence intervals (C.I.) and 95% prediction intervals. If the 95% C.I. did not intersect zero, the results were considered statistically different between the two groups. Positive values of the mean effect sizes indicated that IWT enhanced spermatozoa motility kinetics and enzyme activities compared to the control. Conversely, negative values indicated that these variables were decreased. Furthermore, the Hedges’ g was also interpreted using a rule of thumb similar to the Cohen’s d with some modifications: 0 < |Hedges’ g| ≤ 0.5, 0.5 < |Hedges’ g| ≤ 0.8 and |Hedges’ g| > 0.8 were interpreted as small, medium and large effects, respectively.^39^ Therefore, the actual difference between means of the groups is co-determined by the statistic *P*-value and effect size. The *I*^2^ statistic was used to calculate the proportion of total variation in the estimates of treatment effects owing to inconsistencies in the population effect across studies.^40^ The heterogeneity was interpreted as low, moderate and high when *I*^2^ was 25%, 50% and 75%, respectively. Moderator tests were carried out in the analysis of the dependent variables investigated.

#### Publication bias and sensitivity analysis

Publication bias occurs when the publication process systematically results in studies with specific outcomes being under or overrepresented relative to the totality of studies conducted. To assess publication bias, a funnel plot is commonly used that shows the magnitude and direction of each effect size along with Egger’s regression test. The funnel plot is asymmetric if the *P*-value is <0.05 indicating the existence of the potential publication bias. In this situation, a nonparametric ‘trim-and-fill’ method is needed and an assessment made to ascertain if additional studies are required to mitigate publication bias.^41^

Outliers and influential observations among the estimated measures across studies were identified by Cook’s distance. Additionally, we used influence and leave-one-out diagnostics^42^ as a combined strategy to assess the robustness and reliability of our meta-analysis. After identifying significant outliers through sensitivity analysis, we re-evaluated the pooled effect sizes by excluding the outlier observations to determine the ultimate effect.

#### Moderator analysis

It is necessary to conduct moderator (or subgroup) analyses when a test for heterogeneity yields a significant result (*I*^2^ > 50%). By pooling empirical studies, it became evident that fish species, temperature, and time of post-sperm activation had a significant influence on the primary response variables (MOT, DSM, VCL, VSL and VAP), and were therefore used in moderator analyses. In addition, moderator analyses of enzyme types were also performed for EAES and ANEA.

## 3. RESULTS

Twenty studies were included in the present study (Figure 1) that covered 20 fish species including 11 cold-water fishes and nine warm-water fishes (Figure 2). In total, 63 figures (including line charts and histograms) and six tables were extracted from these studies (Supplementary S2). The studies involved in the present review were Billard and Cosson (1992), Vladiĉ and Jätrvi (1997), Jezierska and Witeska (1999), Williot et al. (2000), Romagosa et al. (2010), Purchase et al. (2010), Huang et al. (2011), Lahnsteiner (2011); Lahnsteiner and Caberlotto (2012), Lahnsteiner and Mansour (2012), Lahnsteiner (2012), Bombardelli et al. (2013), Effer et al. (2013), Le and Pham (2017), Dumorné et al. (2018), Dadras et al. (2019), Dzyuba et al. (2019), Castro et al. (2020), Le et al. (2021) and Rahi et al. (2021).^25,27,30–31,35,43–57^, For cold-water species, the overall effect size was 105 including *k* = 35 (for MOT), 11 (for DSM), 7 (for VCL), 8 (for VAP), 2 (for LIN), 20 (for EAES) and 22 (for ANEA). For warm-water species, the overall effect size was 112 including *k* = 32 (for MOT), 17 (for DSM), 17 (for VCL), 19 (for VSL), 3 (for VAP), 24 (for LIN) (Tables S1-S7).

### 3.1. Summary of overall effect

Based on the overall effect size combining all studies, IWT decreased MOT, and its effect was larger in cold-water species (SMD = -1.22, *P* = 0.0173, *k* = 35) than warm-water fishes (SMD = - 0.95, *P* < 0.0001, *k* = 32) (Table S1, Figure 3a,d). The IWT also showed a large negative effect on DSM in both cold-water species (SMD = -3.18, *P* < 0.0001, *k* = 11) and warm-water species (SMD = -4.20, *P* = 0.0018, *k* = 17) (Table S2, Figure 4a,d). Among spermatozoa velocity parameters, VAP was negatively affected by IWT in cold-water species (SMD = -0.66, *P* = 0.0094, *k* = 8) and warm-water species (SMD = -3.11, *P* = 0.3022, *k* = 3) (Table S5, Figure 5e,f). Also, IWT showed negative effects on EAES in cold-water species (SMD = -4.07, *P* < 0.0001, *k* = 12) but it was without overall effects on ANEA (SMD = 0.10, *P* = 0.8272, *k* = 10) (Table S7, Figure 6a, Figure 7a). In contrast, IWT showed positive effects on VCL (SMD = 1.25, *P* < 0.0001, *k* = 17) and VSL (SMD = 1.50, *P* < 0.0001, *k* = 19) in warm-water species (Tables S3 and S4, Figure 5c,d). However, IWT was without effects on VCL in cold-water species (SMD = 0.75, *P* = 0.7215, *k* = 7) (Table S3, Figure 5b), and on LIN in both cold-water species (SMD = - 0.14, *P* = 0.6519, *k* = 2) and warm-water species (SMD = -0.14, *P* = 0.2184, *k* = 24) (Table S6), and on ANEA in cold-water species (SMD = 0.10, *P* = 0.8272, *k* = 10) (Table S7, Figure 7a).

**Figure 3.**
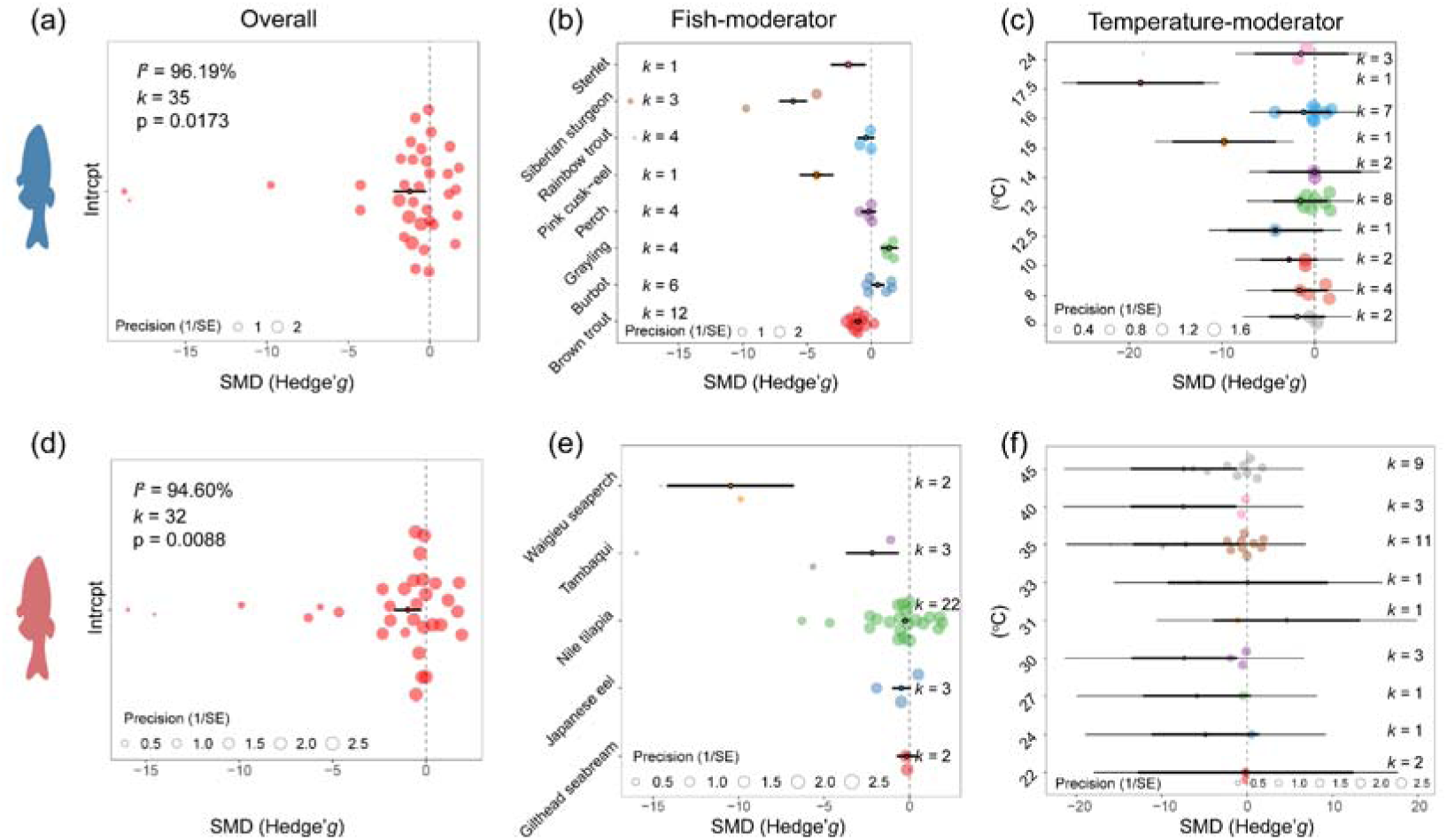
Orchard plots of temperature rise in activation solution showing the impact on sperm motility (%, MOT). The overall effect (**a,d**), as well as fish species-(**b,e**), and temperature-moderator analyses (**c,f**) were illustrated (above: cold fishes-in blue colour, below: warm fishes-in red colour). There are eight, and 10 categories for fish-, and temperature-moderators in cold fishes; five and nine for warm-fishes, respectively. *k* denotes the number of effect sizes for each category of different moderators. For example, *k* = 1, 3, 4, 4, 6 and 12 indicates that 1, 3, 4, 4, 6 and 12 effect sizes were calculated for sterlet (*Acipenser ruthenus*), Siberian sturgeon (*Acipenser baeri*), rainbow trout (*Oncorhynchus mykiss*), grayling (*Thymallus thymallus*), burbot (*Lota lota*) and brown trout (*Salmo trutta*), respectively, when fish species were considered as moderators. SMD is represented by 95% CIs and 95% PIs as scaled effect-size points for each study. Each coloured circle shows a scaled different size of the effect. SMD, standardised mean difference; *I*^2^, the percent variance due to inconsistencies across the studies’ population effect; The statistical significance level was set as *P* < 0.05.

**Figure 4.**
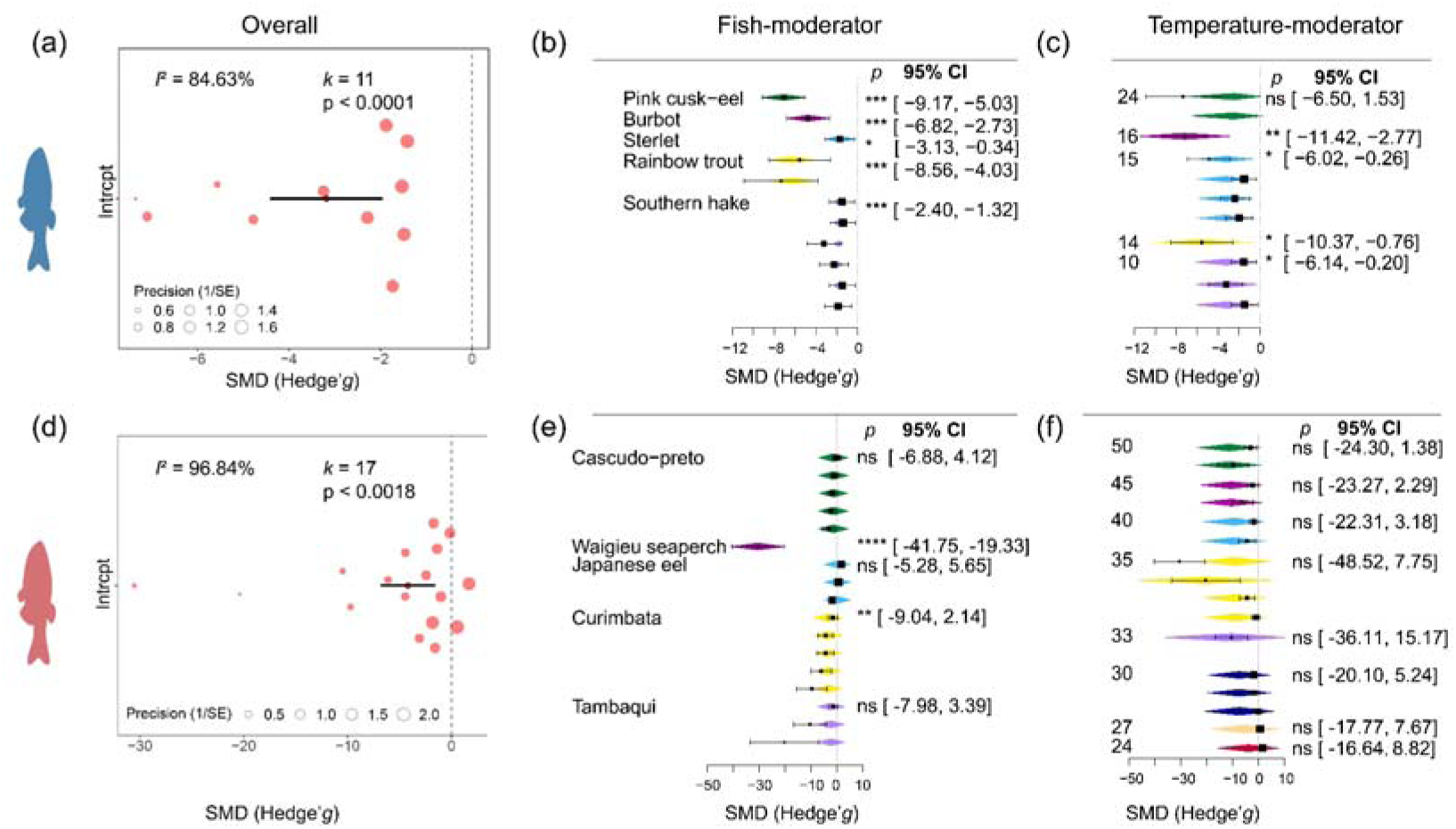
Meta-analytic results of the influence of increased water temperature (IWT) on duration of sperm motility (DSM) of cold-(**a, b** and **c**) and warm-water species (**d, e** and **f**) using the standardised mean difference (SMD, Hedges’g) as the effect size. Orchard plots of IWT impact on DSM in an overall effect (**a, d**). Forest plots of IWT impacts on DSM. Fish species (**b, e**), and temperature (**c, f**) were added as moderators, respectively. The black box set of each forest plot shows the overall effect without any moderators, while the coloured diamond set shows the effect after moderators are added to the model. The categories of each moderator included in this meta-analysis are shown to the left of the forest plots, with different colours representing different categories. SMD is represented by 95% CIs and 95% PIs as scaled effect-size points for each study. Each coloured circle shows a scaled different size of the effect. SMD, standardised mean difference; *I*^2^, the percent variance due to inconsistencies across the studies’ population effect. K denotes the number of effect sizes for each category of different moderators.

**Figure 5.**
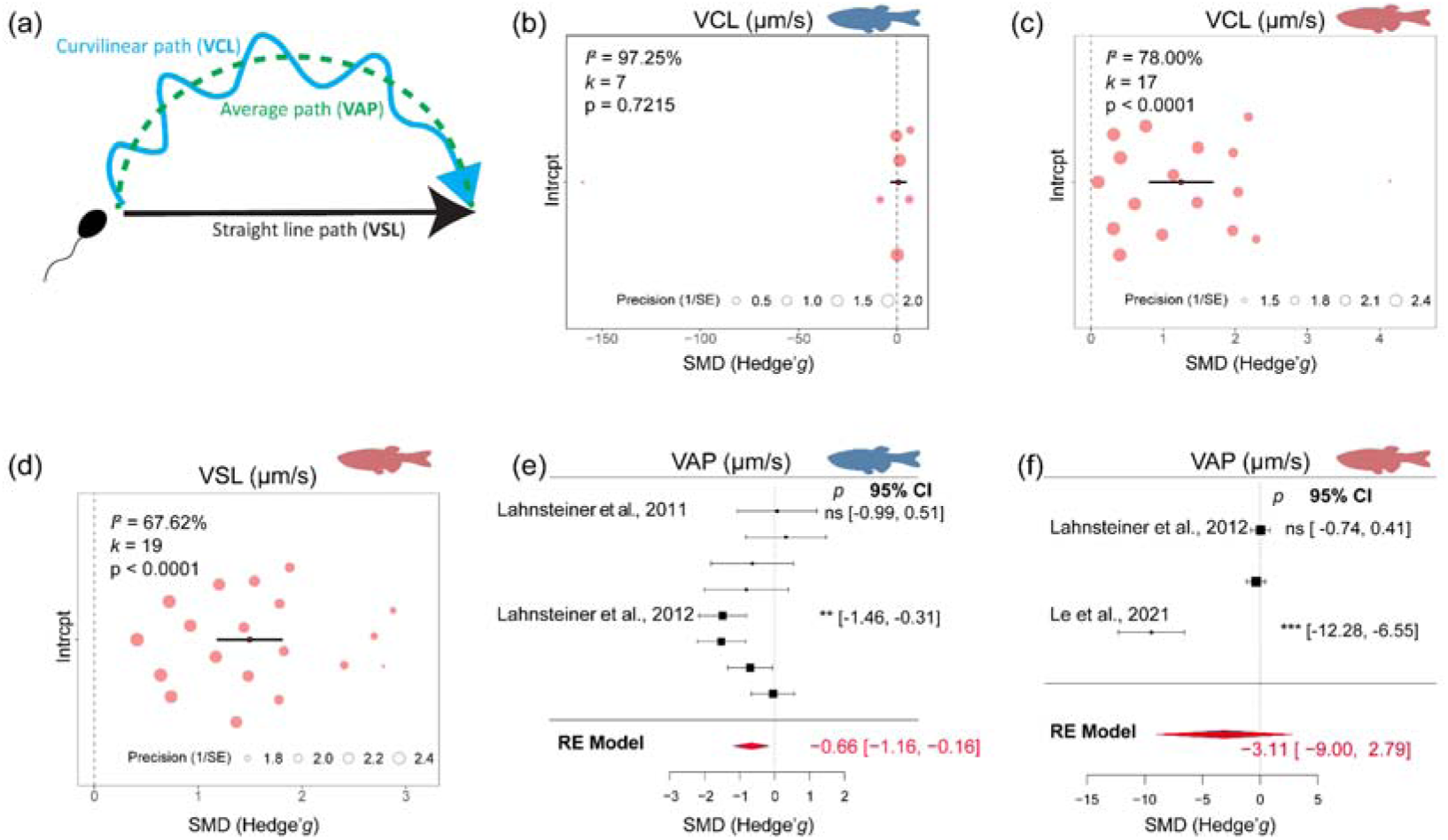
Track of sperm movement showing the different characteristics of the movement as assessed by CASA (VCL, curvilinear velocity; VAP, average path velocity; VSL, straight line velocity) (**a**). Orchard plots of increased water temperature (IWT) overall effect on VCL in cold-(**b**) and warm-(**c**) water fishes, as well as VSL in warm-water fishes (**d**). Forest plots of IWT impact on VAP in cold-(**e**) and warm-water species (**f**). SMD is represented by 95% CIs and 95% PIs as scaled effect-size points for each study. Each colourful circle shows a scaled different size of the effect. SMD, standardised mean difference; *I*^2^, the percent variance due to inconsistencies across the studies’ population effect; K denotes the number of effect sizes for each category of different moderators.

**Figure 6.**
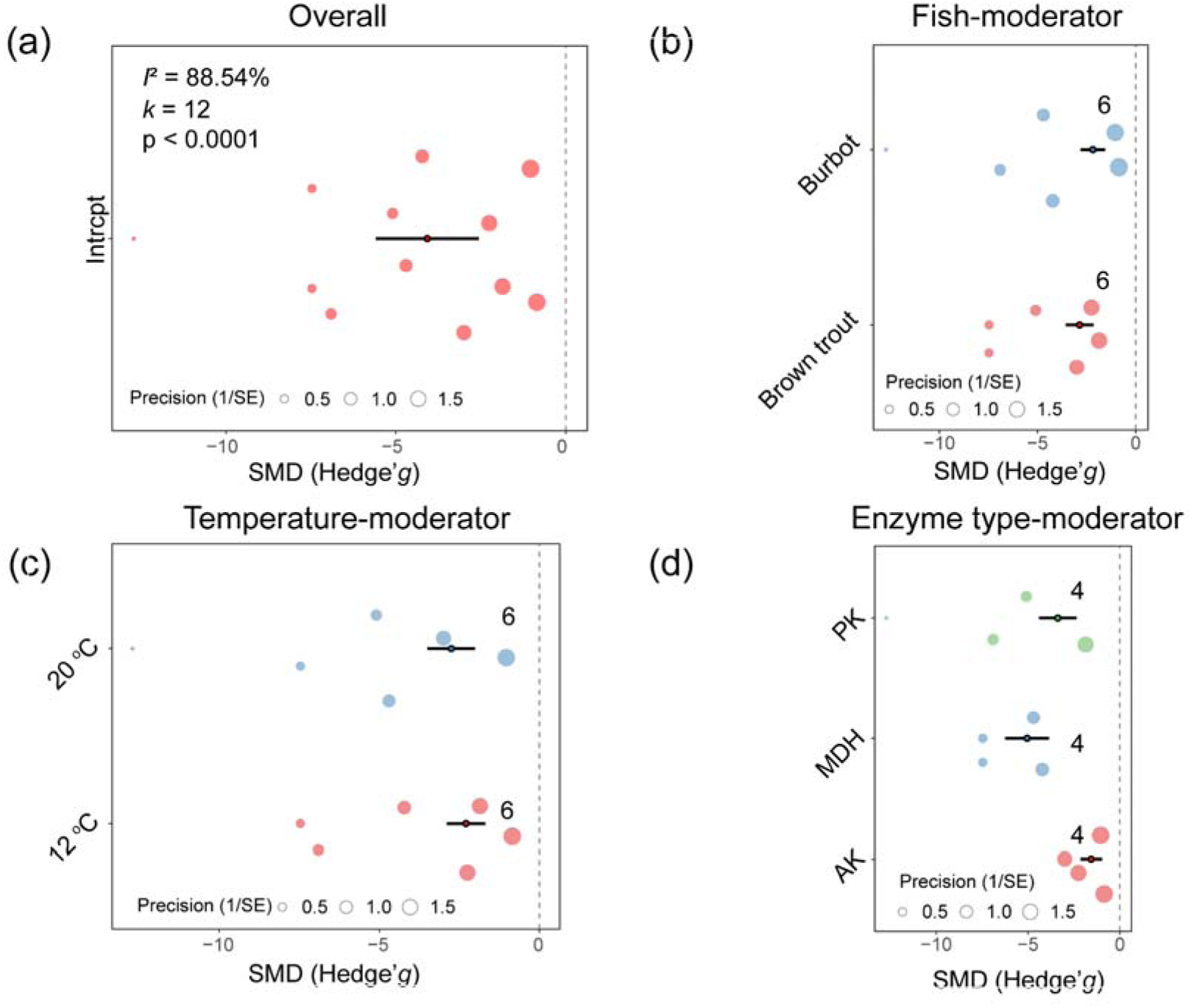
Orchard plots of increased water temperature (IWT) impact on activities of enzymes for sperm energy supply (EAES) in cold water species and overall effect analysis (a), fish- (b), temperature- (c) and enzyme type- (d) moderator analyses. Here are two, five and two categories for fish-, enzyme type- and temperature-moderators, respectively. The number of effect sizes for each category of different moderators are illustrated. For example, 10 in panel (b) indicates that 10 effect sizes were calculated for burbot (*<u>Lota lota</u>*) and brown trout (*Salmo trutta*), respectively, when fish species were regarded as the moderator. SMD is represented by 95% CIs and 95% PIs as scaled effect-size points for each study. Each colourful circle shows a scaled different size of the effect. SMD, standardised mean difference; *I*^2^, the percent variance due to inconsistencies across the studies’ population effect; K denotes the number of effect sizes for each category of different moderators.

**Figure 7.**
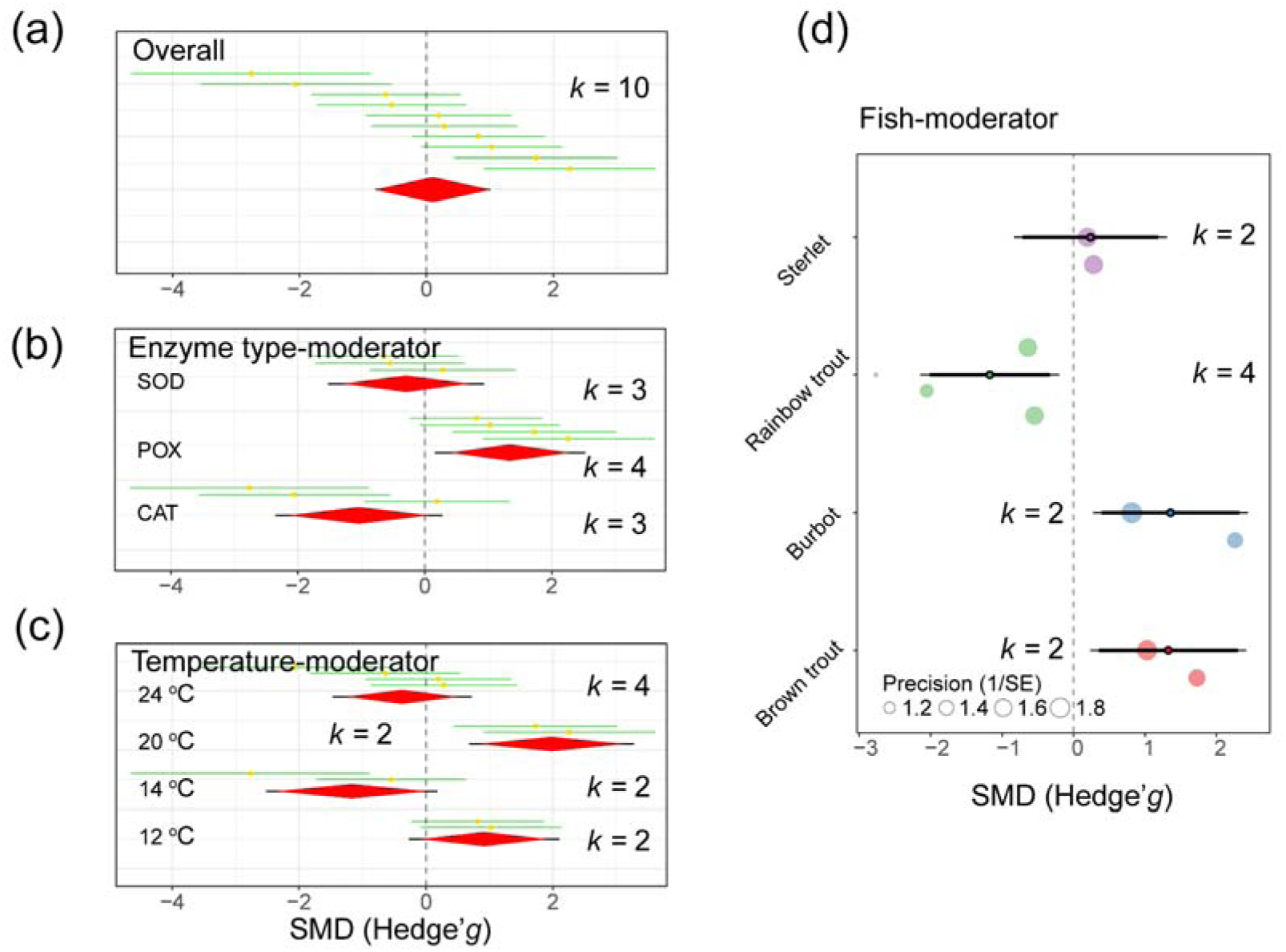
Caterpillar plot illustrating the effect of increased water temperature (IWT) on sperm antioxidant enzyme activity (ANEA) in cold water species using Hedges’ g as the effect size. ANEA underwent overall effect and fish/temperature-moderator analyses. Each green line shows an effect size with a yellow dot (mean effect size). The overall effect size is presented by the red diamond, which includes the mean (centre), the 95% confidence interval (CI) (left and right borders), and the 95% prediction interval (PI) (black line through the diamond). SMD, standardised mean difference; *k*, number of effect sizes for each category of different moderators.

The articles selected reported high heterogeneity for MOT (*I*^2^ = 96.19% and 94.46% for cold- and warm-water species), DSM (*I*^2^ = 84.63% and 96.84%), VCL (*I*^2^ = 97.25% and 78.00%). VAP (*I*^2^ = 63.60% and 98.85%) in cold- and warm-water species, respectively, as well as VSL (*I*^2^ = 59.86%) in warm-water species and EAES (*I*^2^ = 88.54%) and ANEA (*I*^2^ = 80.91%) in cold-water species. However, low heterogeneity was observed for LIN (*I*^2^ = 0.00% and 53.50% in cold- and warm-water species).

These results showed that IWT overall generally decreased MOT, DSM and VAP, but VCL and VSL were increased compared to activated spermatozoa motility at spawning temperatures. However, IWT showed no effect on LIN. In cold-water species, the negative effect of IWT on EAES was observed, while ANEA was not affected.

### 3.2. Moderators affect changes in sperm kinetic parameters

There were significant differences in the effect sizes of spermatozoa motility kinetics after activation at elevated temperatures when fish species, TPSA, temperature and enzyme type were considered as moderators.

#### 3.2.1. IWT decreases percentage of spermatozoa motility (MOT)

When fish species were included in the moderator analyses, overall effects of IWT on MOT showed negative effects in both cold-water and warm-water species (Figure 3a and 3d). Among the species studied, a few species existed in which IWT showed positive effects on MOT, including burbot and grayling (Table S1; Figure 3b).

In cold-water species, IWT effects on MOT were negative in brown trout (SMD = -1.04, *P* < 0.0001, *k* = 4), pink cusk-eel (*Genypterus blacodes*) (SMD = -4.26, *P* < 0.0001, *k* = 1), Siberian sturgeon (*Acipenser baeri*) (SMD = -6.10, *P* < 0.0001, *k* = 3) and sterlet (*Acipenser ruthenus*) (SMD = -1.78, *P* = 0.0107, *k* = 1), and slightly positive in burbot (SMD = 0.51, *P* = 0.0466, *k* = 6) and grayling (SMD = 1.42, *P* < 0.0001 , *k* = 4). However, it was without effects on MOT in perch (*Perca fluviatilis*) (SMD = -0.23, *P* = 0.4513, *k* = 4) and rainbow trout (*Oncorhynchus mykiss*) (SMD = -0.41, *P* = 0.2310, *k* = 4) (Figure 3b).

In warm-water species, IWT negatively affected MOT in Nile tilapia (*Oreochromis niloticus*) (SMD = -0.26, *P*= 0.0045, *k* = 22), Waigieu seaperch (*Psammoperca waigiensis*) (SMD = -10.47, *P* < 0.0001, *k* = 2) and Tambaqui (*Colossoma macropomum*) (SMD = -2.19, *P* = 0.0056, *k* = 3), but was without effects on MOT in gilthead seabream (*Sparus aurata*) (SMD = -0.18, *P* = 0.5457, *k* = 2) and Japanese eel (*Anguilla japonica*) (SMD = -0.48, *P* = 0.0873, *k* = 3) (Figure 3e).

Temperature-moderator analysis revealed that different temperatures had various effects on MOT in cold-water and warm-water species. Temperatures between 15 to 24°C (|effect size| > 1 or/and *P* < 0.05) largely affected MOT in cold-water species (Figure 3c), but temperatures ranging from 35 to 45 °C (|mean effect| > 5, *P* = 0.06-0.07, *k* = 11, 3 and 9 for 35, 40 and 45 °C, respectively) showed large effects on MOT in warm-water species (Figure 3f).

#### 3.2.2. IWT decreases duration of spermatozoa motility (DSM)

Compared to normal temperatures that trigger spermatozoa motility during natural spawning, IWT decreased DSM with negative effect sizes in both cold-water (Figure 4a) and warm-water fishes (Figure 4d). According to the species-moderator analysis, IWT extensively displayed negative effects on DSM in all cold-water species [SMD = -4.76, *P* < 0.0001, *k* = 1 for burbot; SMD = -7.14, *P*p < 0.0001, *k* = 1 for pink cusk-eel; SMD = -6.47, *P* < 0.0001, *k* = 2 for rainbow trout; SMD = -1.74, *P* = 0.0147, *k* = 1 for sterlet, and SMD = -1.86, *P* < 0.0001, *k* =6 for southern hake (*Merluccius australis*)] (Figure 4b). In warm-water species, IWT effects on DSM were smaller for Japanese eel (SMD = 0.18, *P* = 0.8703, *k* = 3) than Cascudo-preto (*Rhinelepis aspera)* (SMD = -1.38, *P*= 0.2392, *k* = 5) and Curimbata (*Prochilodus lineatus*) (SMD = -3.45, *P* = 0.0070, *k* = 5) and Waigieu seaperch (SMD = -30.54, *P* < 0.0001, *k* = 1) (Figure 4e).

The temperature-moderator analysis showed significant effects on DSM in all temperature ranges for both cold-water (10-24 °C) and warm-water fishes (24-50 °C) (|mean effect| > 2.5 or *P* < 0.05) (Figure 4c and f).

#### 3.2.3. IWT alters spermatozoa velocity

The different characteristics of the sperm movement were shown in Figure 5a. A positive effect of IWT on VCL was observed in warm-water species, but it was without effects on cold-water species (Figure 5b and 5c). The TPSA-moderator analysis showed large effect of IWT on VCL at 60 s TPSA for cold-water species, and at 30 s to 360 s TPSA for warm-water species (mean effect > 0.8, *P* < 0.05) (Table S3). However, fish-moderator analysis showed no effects in cold-water species except for sterlet (SMD = 6.33, *P* = 0.8561, *k* = 1) (Table S3).

Compared to VCL, IWT showed significant positive effects on VSL when either TPSA-moderator or temperature-moderator analyses were performed (mean effect > 0.8, *P* < 0.05) (Figure 5d; Table S4).

Negative effects of IWT on VAP were observed in brown trout (SMD = -0.88, *P* = 0.0025, *k* = 4) and Waigieu seaperch (SMD = -9.42, *P* < 0.0001, *k* = 1), but it was without effects on VAP in perch and gilthead seabream (|effect size| < 0.5 and *P* > 0.5) (Table S5, Figure 5e and 5f).

#### 3.2.4. IWT decreases enzymatic activities for energy supply (EAES)

An overall trend of IWT-induced reduction of EAES was observed in cold-water species (Table S7, Figure 6). In this regard, analysis of moderators exhibited negative effects of IWT on EAES. The species-moderator analysis showed IWT negative effects on EAES in brown trout (SMD = - 2.85, *P* < 0.0001, *k* = 6) and burbot (SMD = -2.18, *P* < 0.0001, *k* = 6) with large effect sizes (Figure 6b).

Of five enzymes involved in the EAES-moderator analysis, except peroxidase, four were negatively affected with large effect sizes ranging from -1.55 to -5.04. Additionally, malate dehydrogenase (MDH) was the most powerful enzyme involved in sperm energy supply with the largest effect sizes, followed by pyruvate kinase (PK) and adenylate kinase (AK) (SMD = -5.04, *P* < 0.0001, *k* = 4 for MDH; SMD = -3.37, *P* < 0.0001, *k* = 4 for PK; SMD = -1.55, *P* < 0.0001, *k* = 4 for AK) (Figure 6c). In addition, the large effect size of temperature-moderator analysis was found at 12 and 20 °C (SMD = -1.13, *P* < 0.0001, *k* = 10 for 12 °C; SMD = -1.20, *P* < 0.0001, k = 10 for 20 °C) (Figure 5d).

#### 3.2.5. Species-specific antioxidant enzyme activity (ANEA)

Fish-moderator indicated species-specific effects of IWT on ANEA among cold-water species. The IWT had detrimental effects on rainbow trout (SMD = -1.17, *P* = 0.0062, *k* = 4) but not on sterlet (SMD= 0.24, *P* = 0.6426, *k* = 2) (Figure 7d, Table S7). The IWT effect on ANEA was positive for brown trout (SMD = 1.32, *P* = 0.0076, *k* = 2) and burbot (SMD = 1.36, *P* = 0.0085, *k* = 2). Also, a positive effect of IWT on ANEA was observed in peroxidases (POX) (SMD = 1.34, *P* = 0.0643, *k* = 3) rather than superoxide dismutase (SOD) (SMD = -0.29, *P* = 0.5619, *k* = 3) and catalase (CAT) (SMD = -1.04, *P* = 0.0643, *k* = 3) (Figure 7b). Additionally, temperature specificity on IWT effects on ANEA was observed according to temperature-moderator analysis (Figure 7c). Findings revealed that 20 °C (SMD = 1.98, *P* = 0.0006, *k* = 2) had a large beneficial impact on ANEA, but the effect was negligible at lower temperature (*P* > 0.05).

### 3.3. Publication bias and sensitivity analysis

In cold-water species, the funnel plots and results of Egger’s regression test demonstrated potential publication bias for the effect sizes of DSM, EAES and ANEA (*P* < 0.05), while this was not observed for MOT, VCL and VAP. In these cases, the ‘trim and fill’ method was used to estimate the number of missing effect sizes or studies needed for the current meta-analysis. The results revealed that 1, 1 and 2 additional effect sizes/studies were needed for DSM, EAES and ANEA, respectively, to add and adjust for publication bias (Figure S1). However, IWT had significant effects on the DSM [adjusted SMD = -2.44, *P* < 0.0001, *k* = 12; fail-safe *n* = 297 (the number of additional ‘negative’ studies)]; EAES (adjusted SMD = -1.16, *P* < 0.0001, *k* = 21; fail-safe *n* =253) and ANEA (adjusted SMD = -0.71, *P* = 0.0013, *k* = 11; fail-safe *n* = 16). The influence analysis showed no potential effect size outliers for VAP, LIN, EAES and ANEA, but 2, 1, and 1 outliers of effect size were presented for MOT, DSM and VCL (Figure S2).

The funnel plots and Egger’s test of warn-water species indicated potential publication bias for the effect sizes of MOT, DSM, VCL, LIN and VSL (*P* < 0.001), but this was not observed for VAP (*P* = 0.0622). After ‘trim and fill’ for these parameters, it was shown that no more additional studies were needed for MOT, DSM, VCL and LIN; but two studies were needed in VSL (Figure S3). Nevertheless, IWT still had a significant effect on the number of VSL (adjusted SMD = 1.41, *P* < 0.0001, *k* = 21; fail-safe *n* = 922). Additionally, there were no potential effect-size outliers for VSL (Figure S3). For parameters with outliers, the leave-one-out analysis confirmed that no studies carried a significant deviation of effect size from the overall level if we removed these outliers (Tables S8–S14 and Tables S9-S20 in warm- and cold-water species, respectively).

## 4. DISCUSSION

The present meta-analysis was established to determine the effects of IWT on spermatozoa motility kinetics, determinants of male fertility, and to better understand the impacts of global warming on fish reproduction. In both cold-water and warm-water fish species, an increasing temperature above the normal spawning temperature had a detrimental effect on spermatozoa motility kinetics and altered enzyme activities. In general, IWT decreased MOT and DSM, altered spermatozoa velocity parameters, and decreased the EAES and ANEA. To achieve an accurate conclusion, we determined potential publication bias using Egger’s regression test for DSM, EAES and ANEA in cold-water species, and for MOT, DSM, VCL, LIN and VSL in warm-water species. However, the ’trim-and-fill’ analysis revealed no missing studies for MOT, DSM, VCL and LIN, confirming the consistency of the conclusions. For DSM, EAES, ANEA and VSL, additional studies were also incorporated to address publication bias, yet the conclusions remained unchanged. To sum up, conclusions of the present meta-analysis were not affected by publication bias. Findings of the present study suggest impacts of global warming on spermatozoa functions in fishes, which should be considered in maintaining sustainable fisheries and aquaculture.

In fishes, tolerance to environmental temperature depends on the developmental stage. Spawning adults and embryos have narrower tolerance ranges than larvae and non-reproductive adults, and therefore more vulnerable to climate warming.^58^ In spawning males, it has been shown that changes in the temperature of the environment in which spermatozoa motility is activated, affect motility kinetics in both cold- and warm-water species.^25,27,53,55,59^ The results of our meta-analysis also confirm the adverse effects of IWT on spermatozoa motility kinetics, particularly MOT and DSM. These suggest that temperatures at which males spawn under natural conditions are the most suitable temperature for spermatozoa motility kinetics through long-term biological evolution.^60^ In contrast to the present results of meta-analysis, Dadras et al.^18^ reported temperature-induced increases in spermatozoa motility characteristics. However, spawning temperature was not taken into account in making the comparison, and moreover, only seven articles were reviewed.

The present study shows stronger inhibitory effects of IWT on MOT in cold-water species than those of warm-water species. This might be due to the wider temperature range that warm-water species can tolerate for body maintenance and reproduction compared to cold-water species.^57,61^ Basically, fish species inhabiting cold water tend to have slower metabolism than those that live in warm water. Consequently, temperature affects spermatozoa metabolism and energetics, which are determinants of motility kinetics.^22,57,62–64^ Considering species as a moderator, variations in IWT effects on MOT in warm-water species were shown. These were more negative in Nile tilapia, Waigieu seaperch and Tambaqui compared to gilthead seabream and Japanese eel. This might be due to species-specific metabolic characteristics of spermatozoa. For instance, gilthead seabream spermatozoa show higher thermal tolerance ranging from 4 to 22 °C due to sustainable ATP and phosphocreatine levels.^31^ The present study estimates that the average MOT was reduced by 1.3% and 3.5% in cold-water and warm-water species, respectively, when IWT increased by 1 °C. Similarly, simple calculations from Lahnsteiner and Mansour^50^ and Lahnsteiner^35^ showed that MOT decreased by 2.27% to 2.41% and 0.12% to 2.68%, respectively, when the sperm of brown trout, burbot, grayling and gilthead seabream were activated in the medium with temperature of 1 °C higher than normal spawning conditions. It is worth noting that many factors including the age of the brood fishes, methods of sperm collection and motility analysis are capable of affecting spermatozoa sensitivity to IWT. The effects of temperature on MOT vary during the period of spermatozoa movement. Our meta-analysis showed that IWT perturbs MOT after 60 and 360 s in cold-water and warm-water species. Considering the time required for successful fertilization after spawning of sperm and oocytes,^65–67^ greater temperature tolerance in the spermatozoa of warm-water species equates to an increased capacity for safe fertilization. In the current study, spawning temperatures for cold-water and warm-water species were considered in the range of 3–15 and 18–30 °C, respectively. Under higher temperatures for both categories, percentage of spermatozoa motility was estimated to be 56.40 ± 22.14% and 59.92 ± 38.69% for cold-water and warm-water species (Table S3).

Another critical indicator of male fertility is DSM, which is usually shorter in freshwater species than in marine species;^68^ DSM is also associated with enzymatic activity for energy sources. Our meta-analysis is consistent with previous studies that IWT reduces DSM^15^ and shows greater negative effects of IWT on DSM in warm-water compared to cold-water species. All studies on cold-water species incorporated to the present meta-analysis have demonstrated that IWT decreases DSM, however variations exist among warm-water species. The possible reason could be due to species differences in DSM under natural condition from seconds to several minutes or experimental replicates.^27,51^

Sperm motility requires energy, which is provided by hydrolysis of ATP. During the period of motility, ADP produced from ATP hydrolysis serves as a substrate for ATP regeneration. There are several enzymes involved in energy metabolism in spermatozoa, including adenylate kinase (AK), pyruvate kinase (PK), malate dehydrogenase (MDH), phosphocreatine (PCr) and creatine kinase (CK).^69,70^ Our meta-analysis showed inhibitory effects of IWT on these enzymes in cold-water species (brown trout and burbot), with a higher value for MDH than for PK and AK.^50^ Compared to cold-water species, there was a lack of information for the effects of IWT on energy metabolism in warm-water fishes.

Another determinant of male fertility is spermatozoa velocity parameters, as a spermatozoon should approach an oocyte within a short time in order to fertilize it.^66^ Overall results of this meta-analysis showed the impacts of IWT on spermatozoa velocity parameters in warm-water species including VCL, VSL and VAP, but not LIN. However, data of spermatozoa velocity were very limited. For instance, VCL and VSL were available for Nile tilapia showing positive effects of IWT.^55^ Also, VAP was available for Waigieu seaperch and gilthead seabream, showing negative effects of IWT.^31,56^ Our meta-analysis suggests species-specificity to environmental temperature in terms of spermatozoa velocity, however a larger set of data is required for a clear conclusion. In this regard, Lahnsteiner and Caberlotto^31^ reported no change in spermatozoa velocity of gilthead seabream within the temperature of 4 to 22 °C. Our meta-analysis showed no impacts of IWT on LIN (VSL/VCL) in both cold-water and warm-water species. However, increasing temperature in each study considered showed a decrease in LIN that suggests preference of spermatozoa to move in a spinning-like trajectory rather than a straighter line trajectory, a physiological phenomenon that is regulated by intracellular calcium ions^71^ or the temperature-sensitive ion channel TRPV4.^72^

Spermatozoa possess an antioxidant defence system that protects them from oxidative stress.^50,73,74^ Superoxide dismutase (SOD), catalase (CAT) and peroxidases (POX) are among components in the antioxidant defence system in spermatozoa that are temperature sensitive. Our meta-analysis showed overall effects of IWT on these antioxidant enzymes, although the effect was neutral, negative and positive for SOD, CAT and POX, respectively. These results are consistent with individual studies of sterlet, rainbow trout, burbot and brown trout incorporated into the present meta-results.^25,50^

## Conclusions and future directions

The present study reveals the impacts of IWT on spermatozoa motility kinetics as well as enzymatic functions involved in energy metabolism and oxidative stress. This supports the evidence that an extreme temperature reduces male fertility associated with an increase in sperm abnormalities.^75^ We observed species differences in terms of IWT effects on spermatozoa motility kinetics, which might be due to their adaptation to specific temperature ranges associated with species differences in spermatozoa energetics and metabolism. The present study reveals higher negative effects of IWT on MOT and DSM, while the spermatozoa velocity of some species was not affected or even received positive IWT impacts.

To better elucidate the impacts of global warming on fish reproduction,

1. In males, future studies should enrich the current dataset for clear understanding of IWT on functional characteristics of spermatozoa using a meta-analysis. In this context, the effects of IWT on the other fertility indicators in male fishes including sperm production, and spermatozoa viability, density, morphology and DNA integrity.
2. In females, a meta-analysis study is required to investigate the effects of IWT on functional quality of the oocytes in cold- and warm-water species.
3. For both males and females, individual species studies are required to obtain data from fishes inhabiting various climate environments including tropical, temperate and polar ecosystems.
4. In both males and females, the epigenetic effects of IWT are largely unknown to elucidate the next generation effects of global warming.
5. In both males and females, the effects of IWT on the metabolomic, transcriptomic and proteomic of spermatozoa and oocytes are needed to characterize the modes of action of IWT on the gametes functions.
6. Cutting-edge “omics” technologies are needed to advance our understanding of the global warming’s impacts on fish reproduction.
7. The use of experimental animal models such zebrafish (*Danio rerio*) or medaka (*Oryzias latipes*)^76,77^ provides us with a very valuable advantage to better understand the impacts of IWT at different stages of development.
8. In addition to IWT, there are several environmental factors that affect fish reproduction including environmental pollution,^78–80^ ocean acidification,^81,82^ oxygen level^83^ and salinity.^84−86^ Understanding their interactions is necessary to better elucidate the impacts of global warming on fish reproduction.

## Supporting information

S1.Paper information and effect size

S2.Extracted data

## AUTHOR CONTRIBUTIONS

**Yu Cheng**: conceptualization; methodology; software; data curation; investigation; formal analysis; visualization; writing – original draft; writing – review and editing; supervision. **Jinhai Wang**: software; data curation; formal analysis; visualization; writing – review and editing. **Sayyed Mohammad Hadi Alavi**: conceptualization; writing – original draft; writing – review and editing. **Songpei Zhang**: writing – review and editing. **Zuzana Linhartová**: writing – review and editing. **Deepali Rahi Roy**: conceptualization; writing – review and editing. **Nururshopa Eskander Shazada**: writing – review and editing. **Borys Dzyuba**: writing – review and editing. **Otomar Linhart**: conceptualization; funding acquisition; supervision; project administration; writing – review and editing.

## DATA AVAILABILITY AND SUPPLEMENTARY INFORMATION

Supplementary data and documents to this article can be found online (https://drive.google.com/drive/folders/1WHWZiRmv4KfTNcWU57xklTnDgnaNawTf?usp=sharing).

## ACKNOWLEDGEMENTS

This study was supported by the Ministry of Education, Youth and Sport of the Czech Republic (LRI CENAKVA, LM2023038), by the Czech Science Foundation (23-06426S to OL) and by the National Agency for Agriculture Research, Czech Republic (QK21010141 to OL).

