## Supplementary Tables-02.24 for "Impacts of global warming on fertility of male fishes: evidence from meta-analysis of temperature effects on spermatozoa motility"

**Table S1.** Overall effect size and moderator analyses of increasing temperature of activation medium for spermatozoa motility (MOT) in cold-water and warm-water fishes. The number of effect sizes were indicated by K, standardized mean difference Hedges’*g* (SMD), the % variance due to inconsistencies between the population effect across the studies (*I*^2^), 95% confidence interval (CI) and 95% predicted interval (PI). Time post-sperm activation (TPSA).

|  |  | **Subgroup** | ***K*** | **SMD** | ***I*^2^ (%)** | **95% CI** | ***P* value** | **95% PI** |
| --- | --- | --- | --- | --- | --- | --- | --- | --- |
| Cold-water fishes | Overall |  | 35 | -1.22 | 96.19 | -2.23 to -0.22 | **0.0173** | -2.23 to -0.22 |
|  | Fish-moderator | Brown trout | 4 | -1.04 |  | -1.32 to -0.76 | **<0.0001** | -1.34 to -0.74 |
|  |  | Burbot | 6 | 0.51 |  | 0.01 to 1.02 | **0.0466** | -0.002 to 1.03 |
|  |  | Grayling | 4 | 1.42 |  | 0.77 to 2.08 | **<0.0001** | 0.76 to 2.09 |
|  |  | Perch | 4 | -0.23 |  | -0.81 to 0.36 | 0.4513 | -0.82 to 0.37 |
|  |  | Pink cusk-eel | 1 | -4.26 |  | -5.58 to -2.94 | **<0.0001** | -5.58 to -2.94 |
|  |  | Rainbow trout | 4 | -0.41 |  | -1.09 to 0.26 | 0.2310 | -1.10 to 0.27 |
|  |  | Siberian sturgeon | 3 | -6.10 |  | -7.20 to -5.00 | **<0.0001** | -7.20 to -4.99 |
|  |  | Sterlet sturgeon | 1 | -1.78 |  | -3.15 to -0.41 | **0.0107** | -3.15 to -0.41 |
|  | TPSA-moderator | 10 | 23 | -1.84 |  | -4.52 to 0.84 | 0.1785 | -8.33 to 4.65 |
|  |  | 20 | 3 | -1.49 |  | -4.22 to 1.24 | 0.2855 | -8.00 to 5.02 |
|  |  | 30 | 4 | -0.73 |  | -3.47 to 2.01 | 0.5996 | -7.25 to 5.78 |
|  |  | 40 | 1 | -1.23 |  | -4.00 to 1.54 | 0.3842 | -7.76 to 5.30 |
|  |  | 60 | 1 | -3.07 |  | -6.11 to -0.03 | **0.0479** | -9.72 to 3.58 |
|  |  | 3600 | 2 | -2.35 |  | -5.23 to 0.52 | 0.1087 | -8.92 to 4.22 |
|  | Temperature-moderator | 6 | 2 | -1.62 |  | -4.92 to 1.16 | 0.2248 | -7.71 to 3.95 |
|  |  | 8 | 4 | -1.88 |  | -4.61 to 1.37 | 0.2885 | -7.42 to 4.18 |
|  |  | 10 | 2 | -2.76 |  | -5.81 to 0.29 | 0.0764 | -8.59 to 3.08 |
|  |  | 12 | 8 | -1.55 |  | -4.49 to 1.39 | 0.3014 | -7.33 to 4.23 |
|  |  | 12.5 | 1 | -4.25 |  | -9.38 to 0.88 | 0.1042 | -11.40 to 2.89 |
|  |  | 14 | 2 | -0.03 |  | -5.07 to 5.01 | 0.9909 | -7.11 to 7.05 |
|  |  | 15 | 1 | -9.76 |  | -15.33 to -4.20 | **0.0006** | -17.23 to -2.30 |
|  |  | 16 | 7 | -1.20 |  | -4.12 to 1.72 | 0.4216 | -6.97 to 4.57 |
|  |  | 17.5 | 1 | -18.76 |  | -25.59 to -11.93 | **<0.0001** | -27.21 to -10.32 |
|  |  | 24 | 3 | -1.47 |  | -6.52 to 3.58 | 0.5680 | -8.56 to 5.62 |
| Warm-water fish | Overall |  | 32 | -0.95 | 94.46 | -1.66 to -0.23 | **<0.0001** | -1.66 to -0.23 |
|  | Fish-moderator | Gilthead seabream | 2 | -0.18 |  | -0.75 to 0.40 | 0.5457 | -0.75 to 0.40 |
|  |  | Japanese eel | 3 | -0.48 |  | -1.03 to 0.07 | 0.0873 | -1.03 to 0.07 |
|  |  | Nile tilapia | 22 | -0.26 |  | -0.44 to -0.08 | **0.0045** | -0.44 to -0.08 |
|  |  | Waigieu seaperch | 2 | -10.47 |  | -14.18 to -6.77 | **<0.0001** | -14.18 to -6.77 |
|  |  | Tambaqui | 3 | -2.19 |  | -3.73 to -0.64 | **0.0056** | -3.73 to -0.64 |
|  | TPSA-moderator | 0 | 3 | -2.19 |  | -13.51 to 9.14 | 0.7051 | -18.12 to 13.75 |
|  |  | 5 | 3 | -0.48 |  | -11.71 to 10.75 | 0.9332 | -16.35 to 15.39 |
|  |  | 10 | 3 | -5.32 |  | -11.40 to 0.77 | 0.0870 | -18.08 to 7.45 |
|  |  | 30 | 6 | -3.45 |  | -9.57 to 2.67 | 0.2696 | -16.23 to 9.33 |
|  |  | 60 | 2 | -2.46 |  | -8.60 to 3.68 | 0.4324 | -15.25 to 10.33 |
|  |  | 120 | 4 | -4.44 |  | -10.57 to 1.69 | 0.1557 | -17.22 to 8.34 |
|  |  | 180 | 2 | -4.02 |  | -10.16 to 2.12 | 0.1991 | -16.81 to 8.76 |
|  |  | 360 | 3 | -5.33 |  | -11.43 to 0.77 | 0.0869 | -18.10 to 7.44 |
|  |  | 540 | 2 | -6.64 |  | -12.81 to -0.47 | **0.0350** | -19.44 to 6.17 |
|  |  | 720 | 2 | -6.21 |  | -12.36 to -0.05 | **0.0480** | -19.00 to 6.59 |
|  |  | 900 | 2 | -3.88 |  | -10.02 to 2.25 | 0.2150 | -16.67 to 8.90 |
|  | Temperature-moderator | 22 | 2 | -0.18 |  | -12.78 to 12.43 | 0.9748 | -17.99 to 17.64 |
|  |  | 24 | 1 | -3.02 |  | -11.19 to 1.44 | 0.3270 | -18.96 to 9.21 |
|  |  | 27 | 1 | -4.02 |  | -12.19 to 0.44 | 0.1921 | -19.96 to 8.21 |
|  |  | 30 | 3 | -5.49 |  | -13.54 to -1.15 | 0.0699 | -21.38 to 6.69 |
|  |  | 31 | 1 | 4.65 |  | -3.97 to 13.26 | 0.2903 | -10.61 to 19.90 |
|  |  | 33 | 1 | 0.09 |  | -9.28 to 9.46 | 0.9844 | -15.60 to 15.79 |
|  |  | 35 | 11 | -5.31 |  | -13.35 to -0.98 | 0.0795 | -21.20 to 6.87 |
|  |  | 40 | 3 | -5.61 |  | -13.67 to -1.26 | 0.0642 | -21.50 to 6.57 |
|  |  | 45 | 9 | -5.56 |  | -13.61 to -1.22 | 0.0665 | -21.45 to 6.62 |

**Table S2.** Overall effect size and moderator analyses of increasing temperature of activation medium for duration of spermatozoa motility (DSM) in cold-water and warm-water fishes. The number of effect sizes were indicated by K, standardized mean difference Hedges’*g* (SMD), the % variance due to inconsistencies between the population effect across the studies (*I*^2^), 95% confidence interval (CI) and 95% predicted interval (PI).

|  |  | **Subgroup** | ***K*** | **SMD** | ***I*^2^ (%)** | **95% CI** | ***P* value** | **95% PI** |
| --- | --- | --- | --- | --- | --- | --- | --- | --- |
| Cold-water fish | Overall |  | 11 | -3.18 | 84.63 | -4.41 to -1.96 | **<0.0001** | -4.41 to -1.96 |
|  | Fish-moderator | Burbot | 1 | -4.78 |  | -6.83 to -2.73 | **<0.0001** | -6.83 to -2.73 |
|  |  | Pink cusk-eel | 1 | -7.10 |  | -9.17 to -5.03 | **<0.0001** | -9.17 to -5.03 |
|  |  | Rainbow trout | 2 | -6.29 |  | -8.56 to -4.03 | **<0.0001** | -8.56 to -4.02 |
|  |  | Sterlet sturgeon | 1 | -1.73 |  | -3.13 to -0.34 | **0.0147** | -3.13 to -0.34 |
|  |  | South hake | 6 | -1.86 |  | -2.40 to -1.32 | **<0.0001** | -2.40 to -1.32 |
|  | Temperature-moderator | 10 | 3 | -3.17 |  | -6.14 to -0.20 | **0.0363** | -8.00 to 1.65 |
|  |  | 14 | 1 | -5.56 |  | -10.37 to -0.76 | **0.0232** | -11.69 to 0.56 |
|  |  | 15 | 4 | -3.14 |  | -6.02 to -0.26 | **0.0326** | -7.91 to 1.63 |
|  |  | 16 | 1 | -7.10 |  | -11.42 to -2.77 | **0.0013** | -12.85 to -1.34 |
|  |  | 24 | 2 | -2.48 |  | -6.50 to 1.53 | 0.2252 | -8.01 to 3.04 |
| Warm-water fish | Overall |  | 17 | -4.20 | 96.84 | -6.84 to -1.56 | **0.0018** | -6.84 to -1.56 |
|  | Fish-moderator | Cascudo-preto | 5 | -1.38 |  | -6.88 to 4.12 | 0.2392 | -9.12 to 6.35 |
|  |  | Curimbata | 5 | -3.45 |  | -9.04 to 2.14 | **0.0070** | -11.25 to 4.35 |
|  |  | Japanese eel | 3 | 0.18 |  | -5.28 to 5.65 | 0.8703 | -7.53 to 7.90 |
|  |  | Waigieu seaperch | 1 | -30.54 |  | -41.75 to -19.33 | **<0.0001** | -43.00 to -18.09 |
|  |  | Tambaqui | 3 | -2.30 |  | -7.98 to 3.39 | 0.4286 | -10.17 to 5.57 |
|  | Temperature-moderator | 24 | 1 | -3.91 |  | -16.75 to 5.37 | 0.5473 | -31.64 to 20.26 |
|  |  | 27 | 1 | -5.05 |  | -17.88 to 4.22 | 0.4365 | -32.77 to 19.11 |
|  |  | 30 | 3 | -7.43 |  | -20.20 to 1.77 | 0.2504 | -35.13 to 16.70 |
|  |  | 31 | 1 | -1.38 |  | -8.52 to 20.14 | 0.9135 | -21.69 to 33.31 |
|  |  | 33 | 1 | -10.47 |  | -18.88 to 12.32 | 0.4235 | -31.46 to 24.90 |
|  |  | 35 | 4 | -9.09 |  | -21.89 to 0.07 | 0.1603 | -36.82 to 15.01 |
|  |  | 40 | 2 | -9.56 |  | -22.42 to -0.28 | 0.1414 | -37.30 to 14.60 |
|  |  | 45 | 2 | -10.49 |  | -23.39 to -1.17 | 0.1077 | -38.25 to 13.69 |
|  |  | 50 | 2 | -11.46 |  | -24.43 to -2.07 | 0.0803 | -39.25 to 12.75 |

**Table S3.** Overall effect size and moderator analyses of increasing temperature of activation medium for curvilinear velocity (VCL) in cold-water and warm-water fishes. The number of effect sizes were indicated by K, standardized mean difference Hedges’*g* (SMD), the % variance due to inconsistencies between the population effect across the studies (*I*^2^), 95% confidence interval (CI) and 95% predicted interval (PI). Time post-sperm activation (TPSA).

|  |  | **Subgroup** | ***K*** | **SMD** | ***I*^2^ (%)** | **95% CI** | ***P* value** | **95% PI** |
| --- | --- | --- | --- | --- | --- | --- | --- | --- |
| Cold-water fish | Overall |  | 7 | 0.75 | 97.25 | -3.39 to 4.90 | 0.7215 | -3.39 to 4.90 |
|  | Fish-moderator | Burbot | 2 | 0.73 |  | -67.61 to 69.07 | 0.9832 | -95.91 to 97.38 |
|  |  | Rainbow trout | 4 | -0.19 |  | -68.53 to 68.16 | 0.9957 | -96.84 to 96.46 |
|  |  | Sterlet sturgeon | 1 | 6.33 |  | -62.09 to 74.75 | 0.8561 | -90.37 to 103.03 |
|  | TPSA-moderator | 10 | 3 | 0.82 |  | -0.96 to 2.60 | 0.3675 | -2.08 to 3.72 |
|  |  | 20 | 3 | -0.80 |  | -2.74 to 1.13 | 0.4168 | -3.80 to 2.20 |
|  |  | 60 | 1 | 7.09 |  | 3.27 to 10.90 | **0.0003** | 2.64 to 11.53 |
|  | Temperature-moderator | 14 | 2 | 0.74 |  | -68.21 to 69.70 | 0.9831 | -96.72 to 98.21 |
|  |  | 15 | 2 | 0.73 |  | -68.16 to 69.62 | 0.9833 | -96.69 to 98.16 |
|  |  | 24 | 3 | 0.37 |  | -68.53 to 69.26 | 0.9917 | -97.06 to 97.79 |
| Warm-water fish | Overall |  | 17 | 1.25 | 78.00 | 0.80 to 1.69 | **<0.0001** | 0.80 to 1.69 |
|  | TPSA-moderator | 30 | 6 | 1.55 |  | 1.16 to 1.94 | <0.0001 | 1.16 to 1.94 |
|  |  | 60 | 2 | 0.85 |  | 0.28 to 1.42 | **0.0036** | 0.28 to 1.42 |
|  |  | 120 | 4 | 1.35 |  | 0.92 to 1.78 | **<0.0001** | 0.92 to 1.78 |
|  |  | 180 | 2 | 0.36 |  | -0.17 to 0.89 | 0.1869 | -0.17 to 0.89 |
|  |  | 360 | 2 | 1.09 |  | 0.51 to 1.66 | **0.0002** | 0.51 to 1.66 |
|  |  | 540 | 1 | 0.10 |  | -0.65 to 0.85 | 0.7951 | -0.65 to 0.85 |
|  | Temperature-moderator | 30 | 2 | 0.67 |  | 0.11 to 1.22 | **0.0181** | 0.11 to 1.22 |
|  |  | 35 | 7 | 0.72 |  | 0.41 to 1.02 | **<0.0001** | 0.41 to 1.02 |
|  |  | 40 | 2 | 1.97 |  | 1.29 to 2.64 | **<0.0001** | 1.29 to 2.64 |
|  |  | 45 | 6 | 1.50 |  | 1.12 to 1.88 | **<0.0001** | 1.12 to 1.88 |

**Table S4** Overall effect size and moderator analyses of increasing temperature of activation medium for straight-line velocity (VSL) in warm-water fishes. The number of effect sizes were indicated by K, standardized mean difference Hedges’*g* (SMD), the % variance due to inconsistencies between the population effect across the studies (*I*^2^), 95% confidence interval (CI) and 95% predicted interval (PI). Time post-sperm activation (TPSA).

|  | |  | | **Subgroup** | ***K*** | **SMD** | ***I*^2^ (%)** | **95% CI** | ***P* value** | **95% PI** |
| --- | --- | --- | --- | --- | --- | --- | --- | --- | --- | --- |
| Warm-water fish | Overall | |  | | 19 | 1.50 | 59.86 | 1.18 to 1.81 | **<0.0001** | 1.18 to 1.81 |
|  | | TPSA-moderator | | 30 | 6 | 1.58 |  | 1.21 to 1.94 | **<0.0001** | 1.21 to 1.94 |
|  | |  | | 60 | 2 | 1.55 |  | 0.92 to 2.17 | **<0.0001** | 0.92 to 2.17 |
|  | |  | | 120 | 4 | 1.18 |  | 0.77 to 1.59 | **<0.0001** | 0.77 to 1.59 |
|  | |  | | 180 | 2 | 1.32 |  | 0.69 to 1.95 | **<0.0001** | 0.69 to 1.95 |
|  | |  | | 360 | 2 | 1.18 |  | 0.58 to 1.78 | **0.0001** | 0.58 to 1.78 |
|  | |  | | 540 | 2 | 1.14 |  | 0.54 to 1.75 | **0.0002** | 0.54 to 1.75 |
|  | |  | | 720 | 1 | 2.88 |  | 1.81 to 3.95 | **<0.0001** | 1.81 to 3.95 |
|  | | Temperature-moderator | | 30 | 2 | 0.82 |  | 0.27 to 1.38 | **0.0038** | 0.27 to 1.38 |
|  | |  | | 35 | 8 | 1.84 |  | 1.51 to 2.17 | **<0.0001** | 1.51 to 2.17 |
|  | |  | | 40 | 2 | 1.36 |  | 0.76 to 1.96 | **<0.0001** | 0.76 to 1.96 |
|  | |  | | 45 | 7 | 1.21 |  | 0.89 to 1.53 | **<0.0001** | 0.89 to 1.53 |

**Table S5**. Overall effect size and moderator analyses of increasing temperature of activation medium for average path velocity (VAP) in cold-water and warm-water fishes. The number of effect sizes were indicated by K, standardized mean difference Hedges’*g* (SMD), the % variance due to inconsistencies between the population effect across the studies (*I*^2^), 95% confidence interval (CI) and 95% predicted interval (PI). Time post-sperm activation (TPSA).

|  |  | **Subgroup** | ***K*** | **SMD** | ***I*^2^ (%)** | **95% CI** | ***P* value** | **95% PI** |
| --- | --- | --- | --- | --- | --- | --- | --- | --- |
| Cold-water fish | **Overall** |  | 8 | -0.66 | 63.60 | -1.16 to -0.16 | **0.0094** | -1.16 to -0.16 |
|  | Fish-moderator | Brown trout | 4 | -0.88 |  | -1.46 to -0.31 | **0.0025** | -1.63 to -0.14 |
|  |  | Perch | 4 | -0.24 |  | -0.99 to 0.51 | 0.5264 | -1.13 to 0.64 |
|  | TPSA-moderator | 10 | 3 | -0.66 |  | -2.31 to 1.00 | 0.4369 | -3.43 to 2.11 |
|  |  | 20 | 1 | -0.76 |  | -2.55 to 1.02 | 0.4030 | -3.61 to 2.09 |
|  |  | 30 | 1 | 0.06 |  | -1.70 to 1.83 | 0.9453 | -2.77 to 2.90 |
|  |  | 40 | 1 | 0.71 |  | -1.05 to 2.48 | 0.4265 | -2.12 to 3.55 |
|  |  | 3600 | 2 | -1.48 |  | -3.33 to 0.37 | 0.1173 | -4.37 to 1.41 |
|  | Temperature-moderator | 12 | 2 | -0.27 |  | -1.09 to 0.55 | 0.5763 | -1.09 to 0.55 |
|  |  | 16 | 2 | -0.21 |  | -1.05 to 0.62 | 0.6597 | -1.05 to 0.62 |
| Warm-water fish | Overall |  | 3 | -3.11 | 98.85 | -9.00 to 2.79 | 0.3022 | -9.00 to 2.79 |
|  | Fish-moderator | Gilthead seabream | 2 | -0.17 |  | -0.74 to 0.41 | 0.5730 | -0.75 to 0.41 |
|  |  | Waigieu seaperch | 1 | -9.42 |  | -12.28 to -6.55 | **<0.0001** | -12.28 to -6.55 |
|  | TPSA-moderator | 10 | 2 | -4.59 |  | -13.84 to 4.66 | 0.3308 | -20.48 to 11.30 |
|  |  | 360 | 1 | -4.96 |  | -14.25 to 4.33 | 0.2951 | -20.87 to 10.95 |
|  | Temperature-moderator | 22 | 2 | -0.17 |  | -0.74 to 0.41 | 0.5730 | -0.75 to 0.41 |
|  |  | 35 | 1 | -9.42 |  | -12.28 to -6.55 | **<0.0001** | -12.28 to -6.55 |

**Table S6.** Overall effect size and moderator analyses of increasing temperature of activation medium for linearity of spermatozoa velocity (LIN) in cold-water and warm-water fishes. The number of effect sizes were indicated by K, standardized mean difference Hedges’*g* (SMD), the % variance due to inconsistencies between the population effect across the studies (*I*^2^), 95% confidence interval (CI) and 95% predicted interval (PI). Time post-sperm activation (TPSA).

|  |  | **Subgroup** | ***K*** | **SMD** | ***I*^2^ (%)** | **95% CI** | ***P* value** | **95% PI** |
| --- | --- | --- | --- | --- | --- | --- | --- | --- |
| Cold-water fish | Overall |  | 2 | -0.14 | 0.00 | -0.77 to 0.48 | 0.6519 | -0.77 to 0.48 |
|  | PACT-moderator | 10 | 1 | -0.16 |  | -1.04 to 0.73 |  | -1.04 to 0.73 |
|  |  | 20 | 1 | -0.13 |  | -1.02 to 0.75 |  | -1.02 to 0.75 |
| Warm-water fish | Overall |  | 24 | -0.14 | 53.50 | -0.37 to 0.09 | 0.2184 | -0.37 to 0.09 |
|  | TPSA-moderator | 30 | 9 | -0.24 |  | -0.50 to 0.01 | 0.0616 | -0.50 to 0.01 |
|  |  | 60 | 2 | 0.14 |  | -0.39 to 0.66 | 0.6139 | -0.39 to 0.66 |
|  |  | 120 | 7 | -0.35 |  | -0.64 to -0.06 | **0.0183** | -0.64 to -0.06 |
|  |  | 180 | 2 | 0.34 |  | -0.21 to 0.88 | 0.2238 | -0.21 to 0.88 |
|  |  | 360 | 2 | 0.27 |  | -0.27 to 0.80 | 0.3269 | -0.27 to 0.80 |
|  |  | 540 | 2 | 0.06 |  | -0.47 to 0.58 | 0.8356 | -0.47 to 0.58 |
|  | Temperature-moderator | 10 | 2 | -1.69 |  | -2.32 to -1.07 | **<0.0001** | -2.32 to -1.07 |
|  |  | 15 | 2 | -1.00 |  | -1.57 to -0.44 | **0.0005** | -1.57 to -0.44 |
|  |  | 20 | 2 | -0.23 |  | -0.76 to 0.30 | 0.3963 | -0.76 to 0.30 |
|  |  | 30 | 2 | 0.34 |  | -0.19 to 0.87 | 0.2120 | -0.19 to 0.87 |
|  |  | 35 | 7 | 0.03 |  | -0.25 to 0.31 | 0.8296 | -0.25 to 0.31 |
|  |  | 40 | 2 | -0.22 |  | -0.75 to 0.31 | 0.4160 | -0.75 to 0.31 |
|  |  | 45 | 7 | 0.20 |  | -0.09 to 0.48 | 0.1736 | -0.09 to 0.48 |

**Table S7** Overall effect size and moderator analyses of increasing temperature of activation medium for enzymatic activities of spermatozoa for energy supply (EAES) and antioxidant activity (ANEA) in cold-water fishes. The number of effect sizes were indicated by K, standardized mean difference Hedges’*g* (SMD), the % variance due to inconsistencies between the population effect across the studies (*I*^2^), 95% confidence interval (CI) and 95% predicted interval (PI). Time post sperm activation (TPSA), adenylate kinase (AK), malate dehydrogenase (MDH), pyruvate kinase (PK), catalase (CAT), POX, peroxidases (POX), Superoxide Dismutase (SOD)

|  |  | **Subgroup** | ***K*** | **SMD** | ***I*^2^ (%)** | **95% CI** | ***P* value** | **95% PI** |
| --- | --- | --- | --- | --- | --- | --- | --- | --- |
| EAES | Overall |  | 12 | -4.07 | 88.54 | -5.59 to -2.56 | **< 0.0001** | -5.59 to -2.56 |
|  | Fish-moderator | Brown trout | 6 | -2.85 |  | -3.56 to -2.15 | **< 0.0001** | -3.56 to -2.15 |
|  |  | Burbot | 6 | -2.18 |  | -2.81 to -1.55 | **< 0.0001** | -2.81 to -1.55 |
|  | Enzyme type-moderator | AK | 4 | -1.55 |  | -2.14 to -0.96 | **< 0.0001** | -2.14 to -0.96 |
|  |  | MDH | 4 | -5.04 |  | -6.24 to -3.85 | **< 0.0001** | -6.24 to -3.85 |
|  |  | PK | 4 | -3.37 |  | -4.39 to -2.35 | **< 0.0001** | -4.39 to -2.35 |
|  | Temperature-moderator | 12 | 6 | -2.29 |  | -2.90 to -1.69 | **< 0.0001** | -2.90 to -1.69 |
|  |  | 20 | 6 | -2.75 |  | -3.50 to -2.01 | **< 0.0001** | -3.50 to -2.01 |
| ANEA | Overall |  | 10 | 0.10 | 80.91 | -0.80 to 1.00 | 0.8272 | -0.80 to 1.00 |
|  | Fish-moderator | Brown trout | 2 | 1.32 |  | 0.35 to 2.30 | **0.0076** | 0.23 to 2.42 |
|  |  | Burbot | 2 | 1.36 |  | 0.39 to 2.32 | **0.0058** | 0.27 to 2.44 |
|  |  | Rainbow trout | 4 | -1.17 |  | -2.01 to -0.33 | **0.0062** | -2.15 to-0.20 |
|  |  | Sterlet | 2 | 0.24 |  | -0.71 to 1.19 | 0.6264 | -0.83 to1.31 |
|  | Enzyme type-moderator | CAT | 3 | -1.04 |  | -2.13 to 0.06 | 0.0643 | -2.35 to 0.28 |
|  |  | POX | 4 | 1.34 |  | 0.40 to 2.28 | **0.0050** | 0.15 to 2.53 |
|  |  | SOD | 3 | -0.29 |  | -1.29 to 0.70 | 0.5619 | -1.52 to 0.94 |
|  | Temperature-moderator | 12 | 2 | 0.91 |  | -0.08 to 1.91 | 0.0717 | -0.27 to 2.10 |
|  |  | 14 | 2 | -1.17 |  | -2.36 to 0.02 | 0.0535 | -2.52 to 0.18 |
|  |  | 20 | 2 | 1.98 |  | 0.85 to 3.11 | **0.0006** | 0.68 to 3.28 |
|  |  | 24 | 4 | -0.38 |  | -1.26 to 0.51 | 0.4048 | -1.47 to 0.72 |

**Table S8 The results of the leave-one-out analysis for percentage of motile sperm (MOT) in cold-water species**

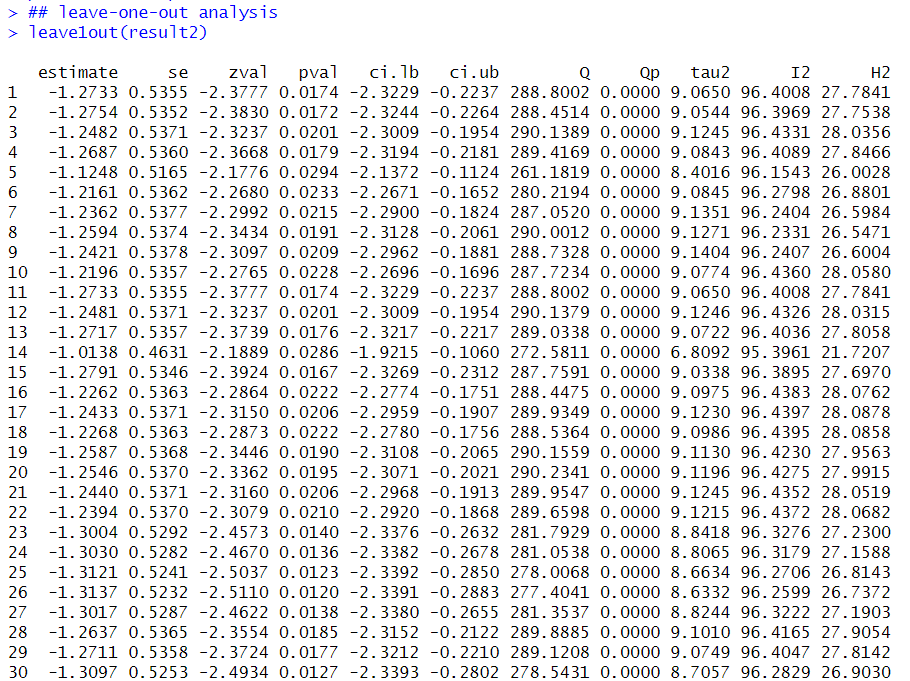

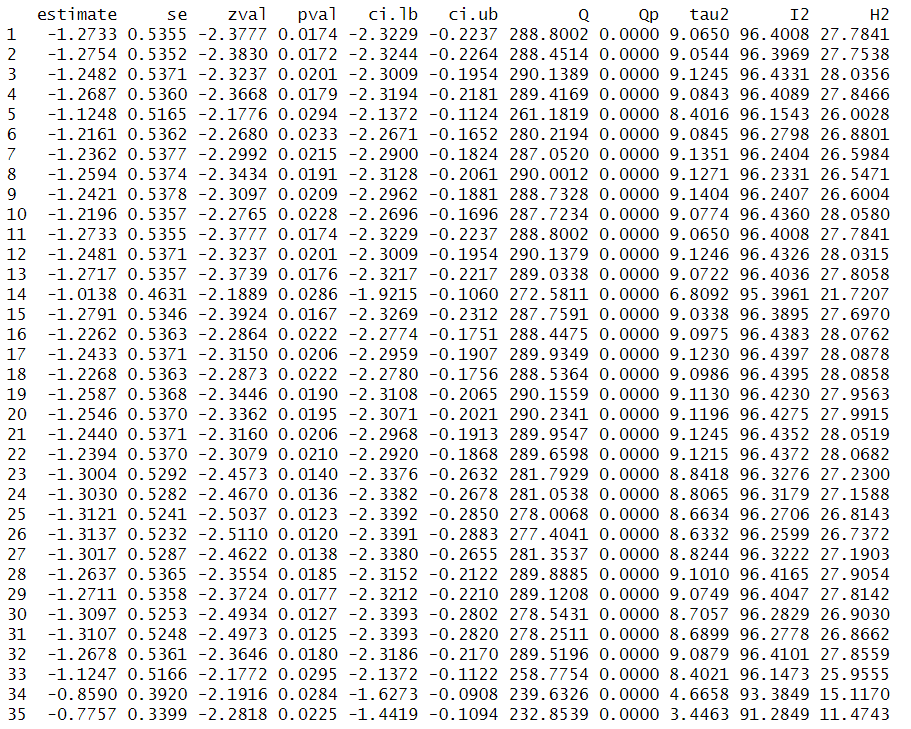

**Table S9 The results of the leave-one-out analysis for duration of sperm motility (DSM) in cold-water species**

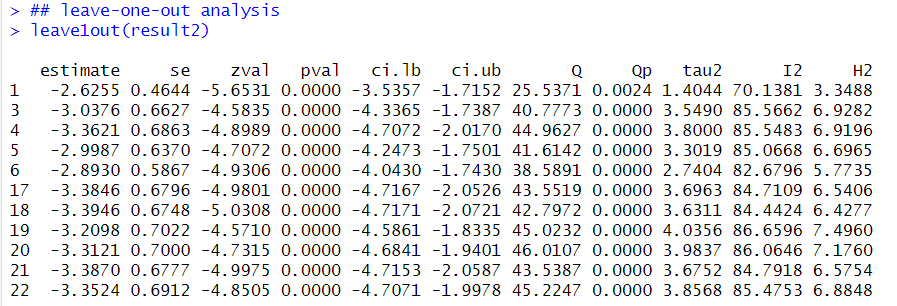

**Table S10 The results of the leave-one-out analysis for sperm VCL in cold-water species**

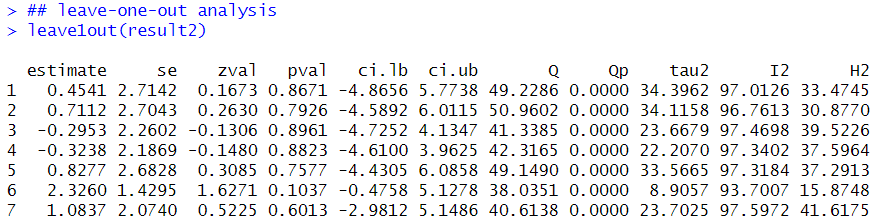

**Table S11 The results of the leave-one-out analysis for sperm VAP in cold-water species**

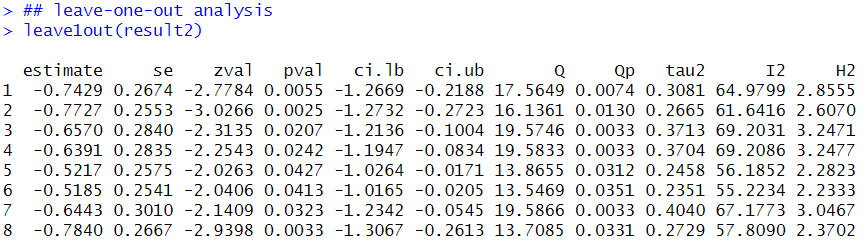

**Table S12 The results of the leave-one-out analysis for sperm LIN in cold-water species**

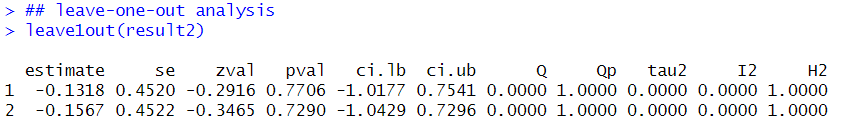

**Table S13 The results of the leave-one-out analysis for activities of enzymes for energy supply (EAES) in cold-water species**

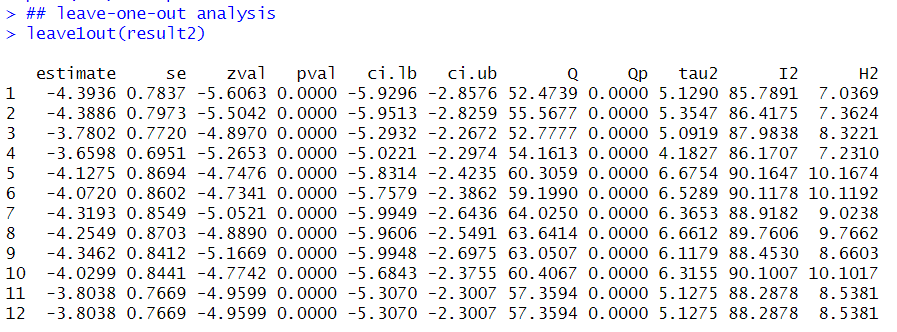

**Table S14 The results of the leave-one-out analysis for antioxidant enzyme activity (ANEA) in cold-water species**

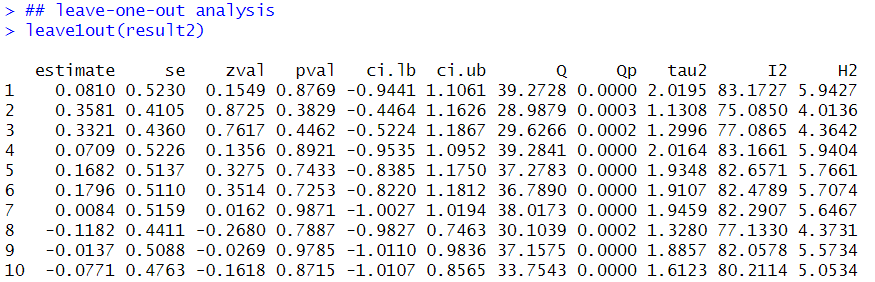

**Table S15 The results of the leave-one-out analysis for percentage of motile sperm (MOT) in warm-water species**

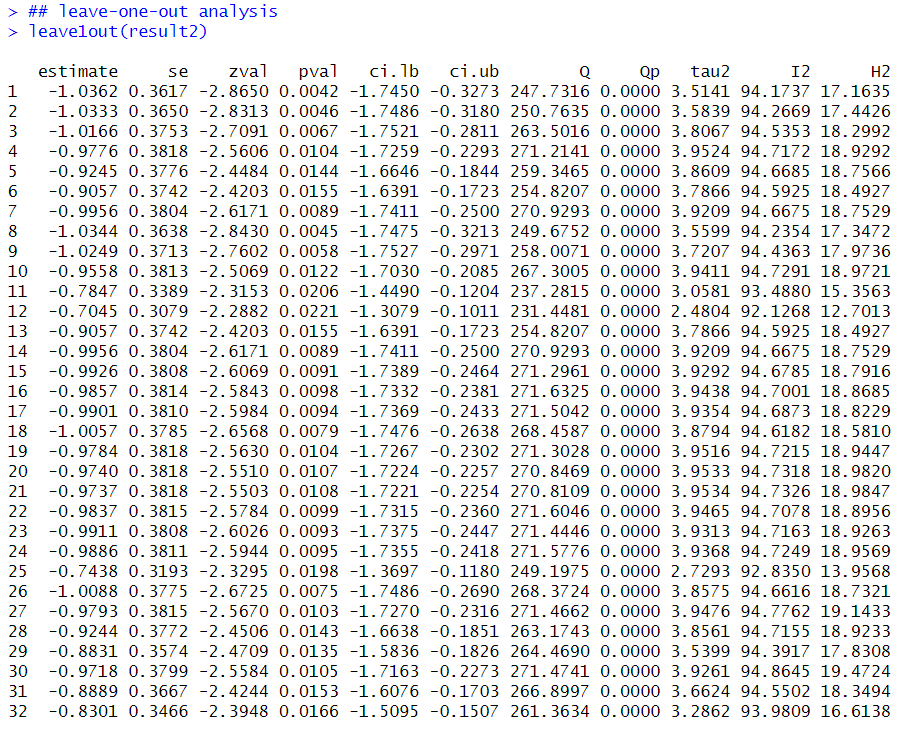

**Table S16 The results of the leave-one-out analysis for duration of sperm motility (DSM) in warm-water species**

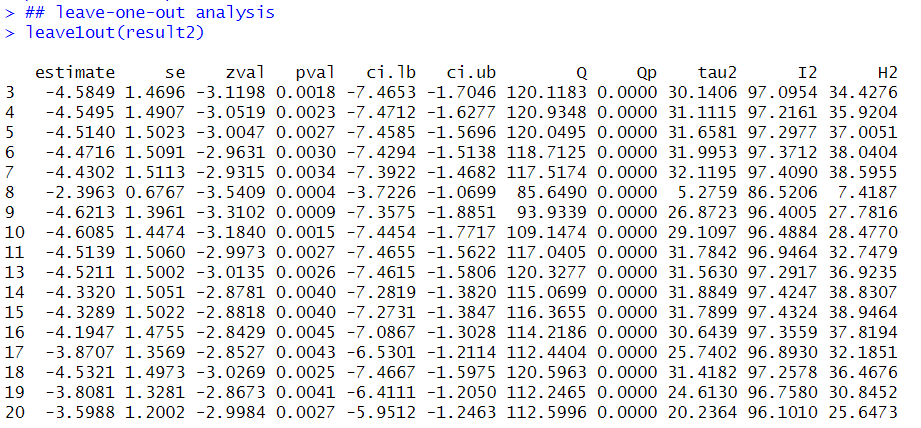

**Table S17 The results of the leave-one-out analysis for sperm VCL in warm-water species**

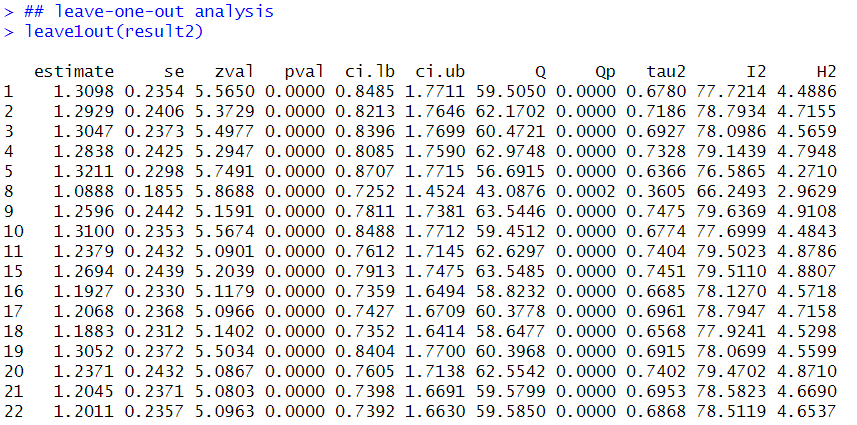

**Table S18 The results of the leave-one-out analysis for sperm VAP in warm-water species**

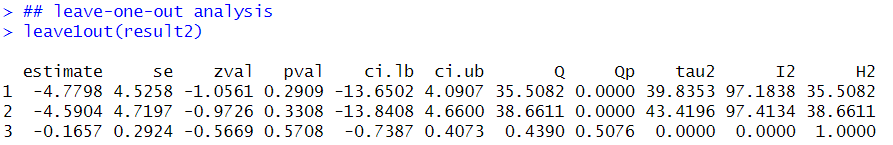

**Table S19 The results of the leave-one-out analysis for sperm LIN in warm-water species**

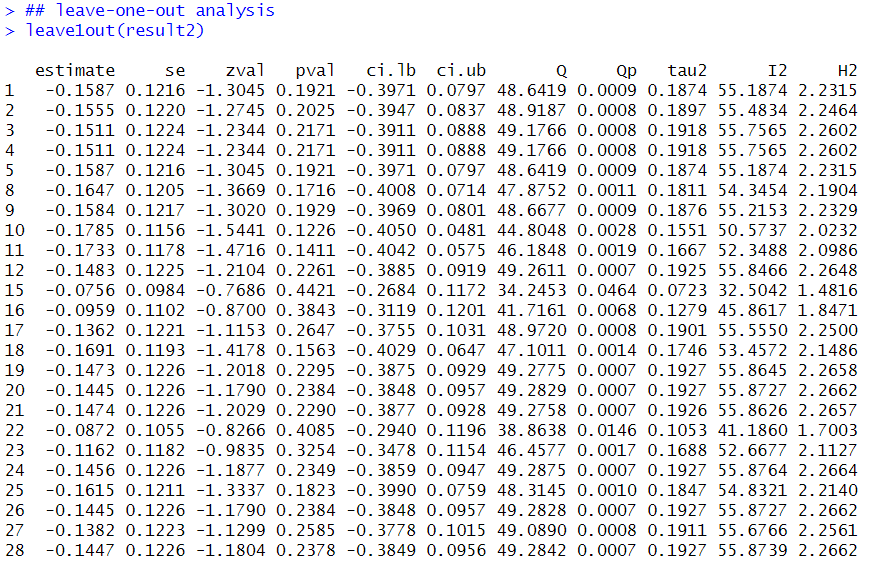

**Table S20 The results of the leave-one-out analysis for sperm VSL in warm-water species**

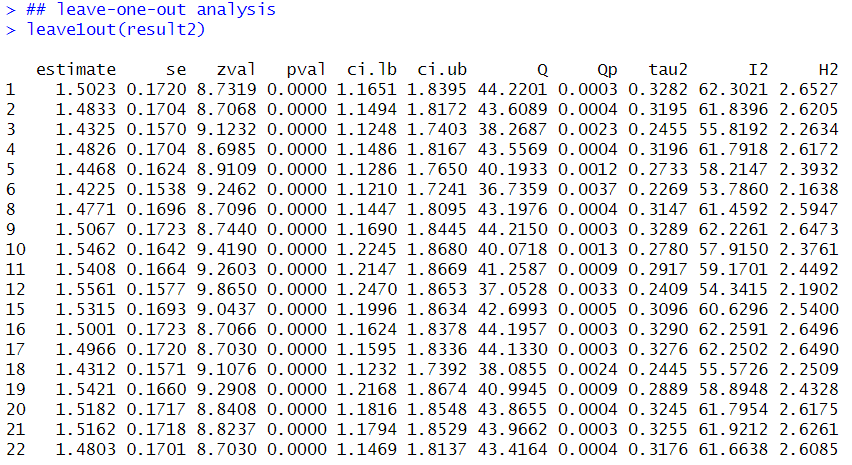
