## Supplement figs for "Impacts of global warming on fertility of male fishes: evidence from meta-analysis of temperature effects on spermatozoa motility"

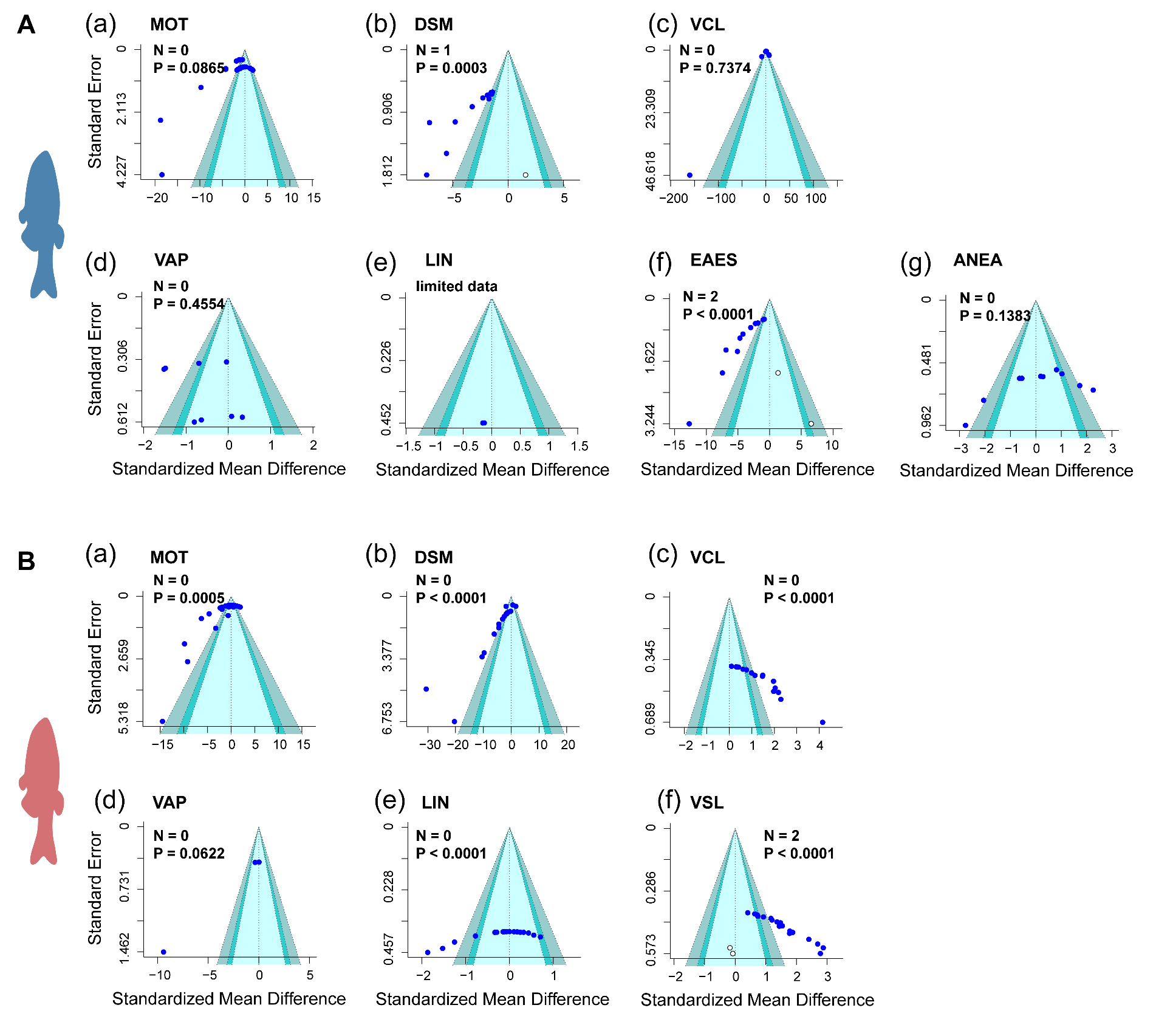

**Figure S1.** Assessment of publication bias. Contour-enhanced funnel plots of standardised mean difference (Hedges' g) for cold-water fishes of percentage of sperm motility (MOT) (A-**a**), duration of sperm motility (DSM) (A-b), sperm velocity (VCL: A-c; VAP: A-d; LIN: A-e), activities of enzymes for energy supply (EAES) (A-f) and antioxidant enzyme activity (ANEA) (A-g) before and after ‘trim and fill’ analysis; warm-water fishes of MOT (B-a), DSM (B-b), VCL (B-c), VAP (B-d), LIN (B-e) and VSL (B-f). Egger's regression test, accompanying *P* values and confidence intervals (CIs) were illustrated in each panel. For EAES, and ANEA in cold-water, and MOT, DSM, VCL, VSL and LIN in warm-water fishes, funnel plots exhibited substantial asymmetry (*P* < 0.05), indicating potential publication bias. After ‘trim and fill’, in cold-water species, 0, 1, 0, 0, 1 and 2 additional studies/effect sizes (white dots) need to be implanted to eliminate publication bias for MOT, DSM, VCL, VAP, LIN, EAES and ANEA, respectively. No study needs to be implanted for MOT, DSM, VCL, VAP and LIN, respectively in warm-water fishes, but two additional studies need to be implanted. 0.05 < *P* < 0.1, 0.01 < *P* < 0.05 and *P* < 0.01 with different colours present 90% CI, 95% CI and 99% CI, respectively. *n*, the number of extra effect sizes/studies that need to be added/deleted, and dark blue dots represent the effect sizes from recruited studies in the current meta-analysis. For example, *n* = 1 for DSM (A-b) suggests that two potential effect sizes/studies need to be added to counteract publication bias. *P* values in the figures indicate statistic results from Egger’s test. Furthermore, one of them were outside the 90% CI region, implying that results of this one investigation were not statistically significant.

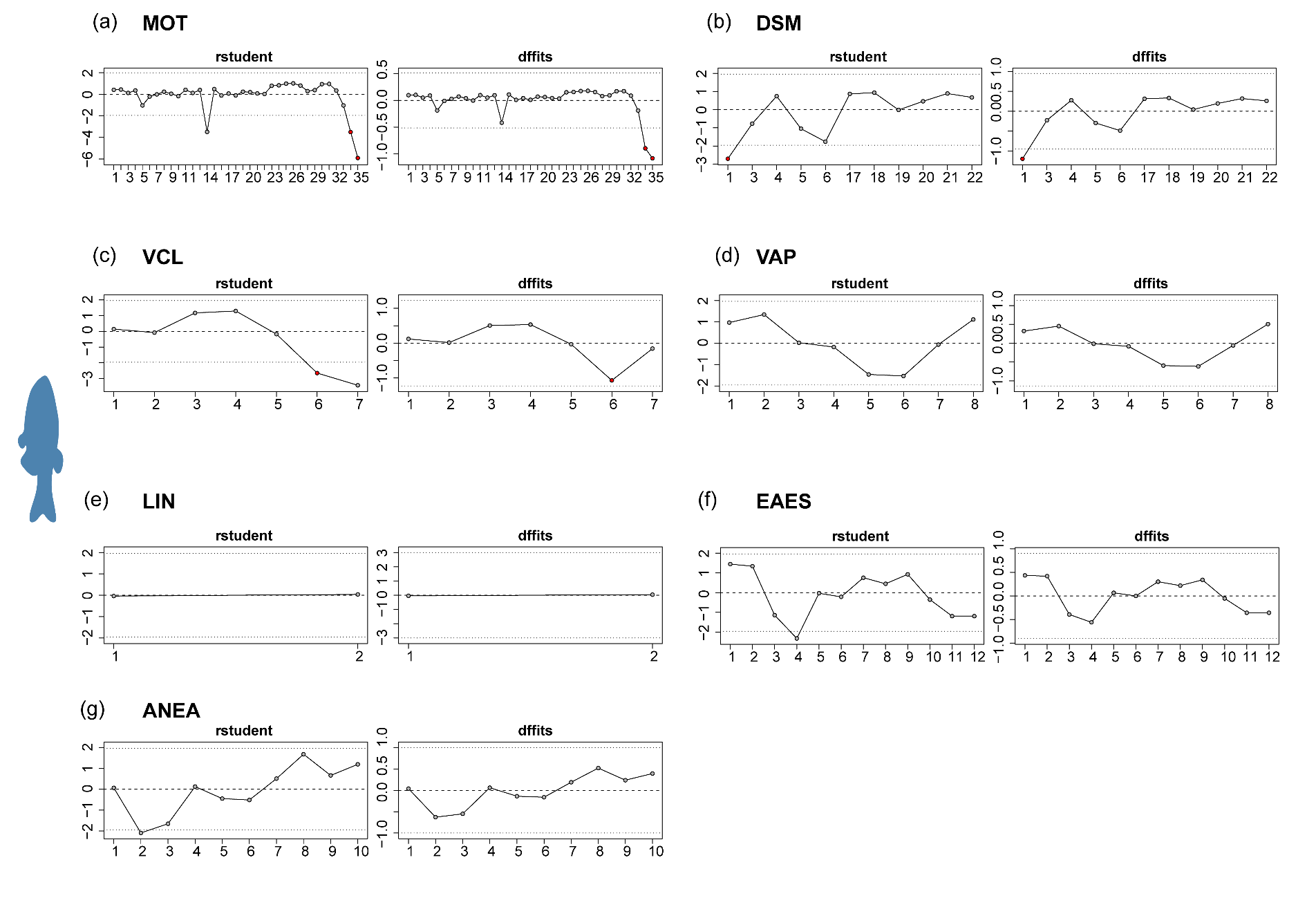

**Figure S2.** Analysis of sensitivity using the influence approach in cold-water species. There are no potential effect-size outliers for average path velocity (VAP) (d), linearity (LIN) (e), activities of enzymes for energy supply (EAES) (f) and antioxidant enzyme activity (ANEA) (g), but 2, 1, 1 outliers of effect size were presented for parameters: percentage of motile sperm (MOT) (a), duration of sperm motility (DSM) (b) and curvilinear velocity (VCL) (c), respectively. The leave-one-out strategy was used to see if these outliers had opposing effects on the outcomes in this situation. Note: outliers are shown by red dots. Please refer to Tables S8 to S14 for the results of the leave-one-out analysis.

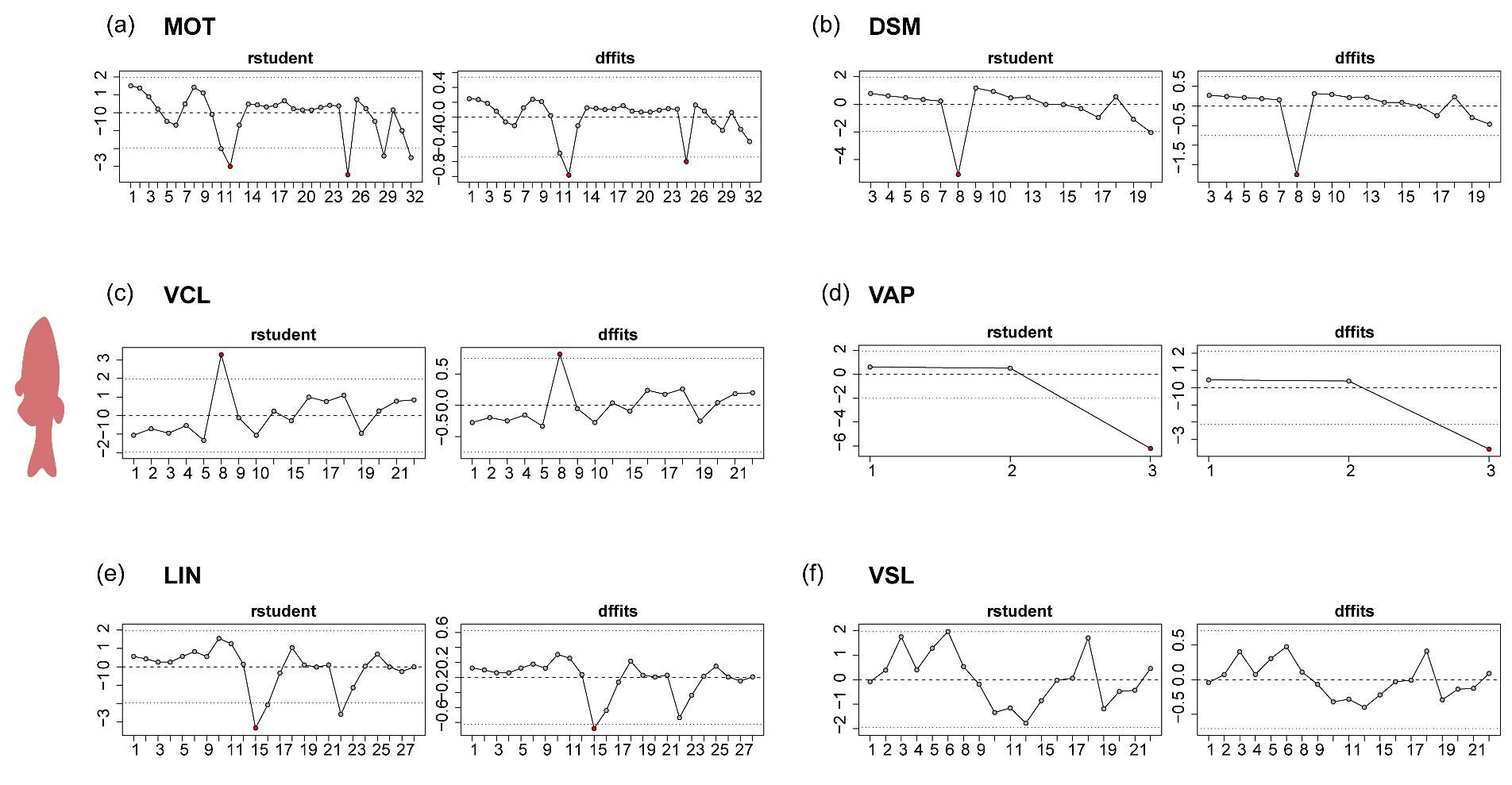

**Figure S3.** Analysis of sensitivity using the influence approach in warm-water species. There are no potential effect-size outliers for (VSL) (f), but 2, 1, 1, 1 and 1 outliers of effect size were presented for parameters: percentage of motile sperm (MOT) (a), duration of sperm motility (DSM) (b), curvilinear velocity (VCL) (c), average path velocity (VAP) (d), Linearity (LIN) (e), respectively. The leave-one-out strategy was used to see if these outliers had opposing effects on the outcomes in this situation. Note: outliers are shown by red dots. Please refer to Tables S15 to S20 for the results of the leave-one-out analysis.
